# An E-I Firing Rate Model for Gamma Oscillations

**DOI:** 10.1101/2023.05.02.539022

**Authors:** Yiqing Lu, John Rinzel

## Abstract

Firing rate models for describing the mean-field activities of neuronal ensembles can be used effectively to study network function and dynamics, including synchronization and rhythmicity of excitatory-inhibitory populations. However, traditional Wilson-Cowan-like models, being amenable to mathematical analysis, are found unable to capture some dynamics such as Interneuronal Network Gamma oscillations (ING) although use of an explicit delay can help. We resolve this issue by introducing a mean-voltage variable (*v*) that considers the subthreshold integration of inputs and works as an effective delay in the negative feedback loop between firing rate (*r*) and synaptic gating of inhibition (*s*). Here we describe a three-variable *r-s-v* firing rate model for I-I networks, which is biophysically interpretable and capable of generating ING-like oscillations. Linear stability analysis, numerical branch-tracking and simulations show that the rate model captures many of the common features of spiking network models for ING. We also propose an alternative *r-u-s* model for ING which considers an implicit delay by a pre-synaptic variable *u*. Furthermore, we extend the framework to E-I networks. With our six-variable *r-s-v* model, we describe for the first time in firing rate models the transition from Pyramidal-Interneuronal Network Gamma (PING) to ING by increasing the external drive to the inhibitory population without adjusting synaptic weights. Having PING and ING available in a single network, without invoking synaptic blockers, is plausible and natural for explaining the emergence and transition of two different types of gamma oscillations.

**Author Summary:** Gamma rhythm (30-90 Hz) is a neuronal oscillation broadly observed across various animal species. It is associated with cognitive functions and gamma oscillation dysfunction has been found in neurological diseases. Traditional studies of gamma oscillations are primarily conducted in detailed but computational demanding spiking network models. Motivated by the growing size of accessible datasets and large-scale modeling, we develop mean-field representations of gamma oscillations that are computationally lightweight and mathematically analyzable. We resolve the incapability of generating oscillation in inhibitory network with traditional Wilson-Cowan firing rate model by introducing implicit delay with a mean membrane potential variable and generalize the model for excitatory-inhibitory network to capture richer dynamics. Our work provides a potential building block for multi-area large-scale modeling to help understanding the functional roles of circuits at different levels.

## Introduction

Gamma oscillations, spanning the range 30-90 Hz [1–3] are found in various brain regions including hippocampus [4–6], entorhinal cortex [7], thalamus [8], visual cortex [9–12] and other neocortical areas [13] etc. Functionally, gamma rhythms are believed to be related to working memory [14,15], attention [16–18], perception [19–21], etc. Abnormal gamma oscillations are seen in cognitive disorders such as Schizophrenia, epilepsy, autism and Alzheimer’s disease [22]. Therefore, studying how gamma oscillations are generated and affected by factors are key to understanding behavior and neurological disease states.

Numerous experimental and theoretical studies consider the mechanisms of gamma oscillation generation, among which two are prominent: Pyramidal-Interneuronal Network Gamma (PING) and Interneuronal Network Gamma (ING). PING is based on the reciprocal interaction of pyramidal cells (excitatory cells, abbreviated as E cells) and fast-spiking PV+ interneurons (inhibitory cells, abbreviated as I cells) [23–25]; ING only involves the recurrent interaction of inhibitory neurons [24,26–28]. There is experimental evidence for both mechanisms. For example, in-vitro experiments in rat CA3 area showed that cholinergically induced gamma oscillation can be blocked by both AMPA and GABA_A_ receptor antagonists [29], suggesting both E and I cells participate in the oscillation; while kainate-induced gamma oscillation can only be blocked by GABA_A_ receptor antagonist but not AMPA receptor antagonist [30], suggesting that I cells alone can support the oscillation.

In theoretical approaches, PING and ING are traditionally studied with spiking network models. The models are detailed, with explicit synaptic kinetics and coupling architecture, but complicated for mathematical analysis and computationally demanding for large scale simulations. Firing rate approaches track the mean firing rate of a population without describing each individual cell’s membrane potential and synaptic activation. Although advantageous computationally and for analytical tractability, the rate framework suffers from incapability to capture certain dynamics, e.g., the kinetics of the inhibitory synaptic field that are important for gamma oscillations. Taking off from the classical Wilson-Cowan model [31], some recent studies have proposed rate models for gamma oscillations. Keeley et al. [32,33] developed Wilson-Cowan like E-I firing rate models (*r-s* model) for PING, by including a synaptic activation variable for each population that can generate robust gamma oscillation emergent via Hopf bifurcation. Such models, however, cannot capture ING dynamics; the network basal state (a fixed point) is stable in the corresponding I-I *r-s* model [34]. Two possible resolutions are: (1) two-variable *r-s* delay model [32], that adds an explicit delay between the firing rate and the synaptic activation; (2) three-variable QIF-FRE model [34], which is an exact macroscopic description of a quadratic integrate-and-fire network. Both models generate ING-like oscillations via Hopf bifurcations, yet the *r-s* delay model is challenging for mathematical analysis due to the explicit delay and the QIF-FRE model is not readily interpretable biophysically from the mean-field equations.

Moreover, PING and ING were usually studied independently – PING in an E-I network and ING in an I-I network, leaving open questions about how ING can dominate in an E-I network without blocking or freezing input from E. Some recent studies with spiking networks looked into the interaction of PING and ING in an E-I network. For example, Börgers & Walker [35], extending an earlier study by Börgers & Kopell [36], showed that with continuously rising drive to I cells, PING can be switched into ING. Viriyopase et al. [37] studied the cooperation and competition of PING and ING and found the mechanism that generate higher oscillation frequency dominates. Nevertheless, little work has been done from the firing rate model perspective.

Therefore, we propose in this paper a biophysically interpretable *r-s-v* firing rate model that captures both PING and ING dynamics. We introduce an additional variable to the *r-s* model [32,33], mean-voltage, *v*(*t*), that represents subthreshold integration of inputs and acts as an effective delay in the negative feedback loop between firing rate, *r*(*t*), and synaptic gating of inhibition *s*(*t*).

Our presentation is organized as follows. In Section 1, we first review PING and ING mechanisms by an example where PING and ING are each realizable in a E-I spiking model with fixed synaptic weights. From simulated network oscillations, we generate across-cell average quantities as mean-field representations to motivate the formulation of our firing rate models. In Section 2, we develop a rate model for ING in an isolated I-I network by introducing the mean membrane potential variable *v*, that provides an implicit delay. We show that our *r-s-v* firing rate model captures many features of an I-I spiking network. We next demonstrate in Section 3 by a combination of numerical and analytical bifurcation analysis that over different parameter variations the model’s behavior matches trends that are consistent with many of those reported in experiments and spiking models. In Section 4, we propose the *r-u-s* model with the variable *u* providing an alternative mechanism for an implicit delay and we compare its behaviors with those of the *r-s-v* model. In Section 5, we extend the *r-s-v* formulation to both subpopulations in an E-I network and show the rate model reproduces a key feature in the spiking network (Section 1) that the network can be tuned from PING to ING by only increasing the input drive to the I population.

## Results

### 1 Coexistence of PING and ING in spiking network models by tuning input drive to the inhibitory cells

Pyramidal-Interneuronal Network Gamma (PING) and Interneuronal Network Gamma (ING) are two major mechanisms that account for the gamma-band oscillation in local field potentials [28,38,39]. Although they are traditionally studied separately in Excitatory-Inhibitory (E-I) and Inhibitory-Inhibitory (I-I) networks, a more natural scenario is that PING and ING are each realizable in the same E-I network and can be activated selectively by inputs or induced by neuromodulation of coupling. Here, we demonstrate the conditional activation by different inputs without altering synaptic coupling. As illustrative, network dynamics can be transitioned from PING (Fig 1A) to ING (Fig 1B) by increasing the extrinsic steady input drive *I*_I_ to the I cells in an E-I spiking network of Type I Hodgkin-Huxley like biophysical neurons [40] with fixed connectivity (see Methods 4.1 for description of cellular and network parameters). The E cells are driven by heterogeneous inputs such that the mean intrinsic frequency is around 40 Hz (see Methods 4.1 for the distribution of *I*_E_ and the exact range of intrinsic frequency), while the I cells are driven by subthreshold or suprathreshold heterogeneous inputs for PING or ING, respectively. To study the dynamical roles of E and I cells in generating the rhythms, we first show raster plots as a general overview for the comparison of firing patterns, synchrony level, cellular and network frequencies. As revealed in numerous experimental and computational studies, the dynamics of inhibition plays a critical role in gamma oscillations – for example, tetanic stimulation-evoked CA1 gamma oscillations can be blocked by GABA_A_ receptor antagonist [3,26,27]. Therefore, we highlight inhibition in the following discussion.

**Fig 1.**
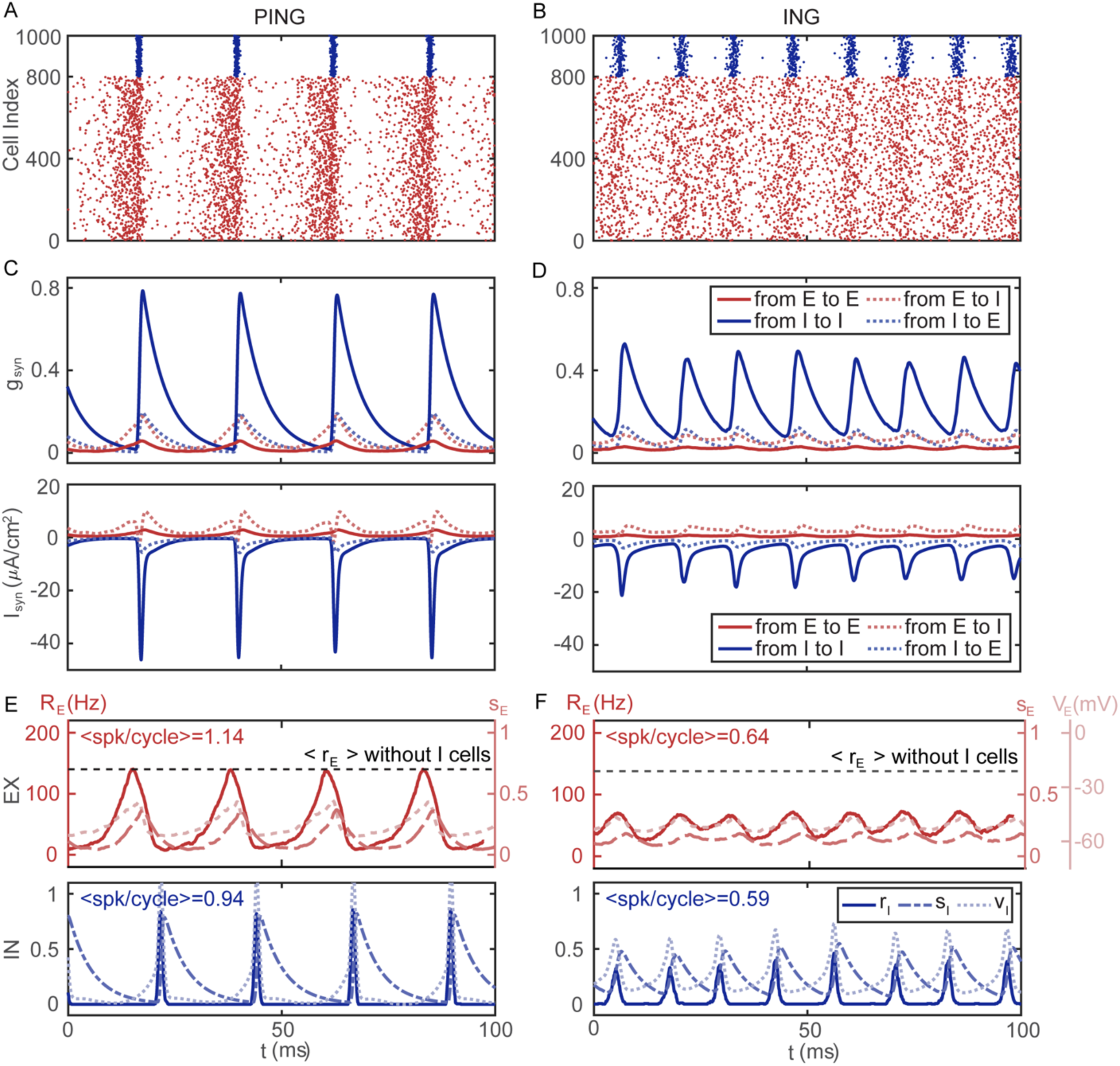
Increasing the drive current *I*_I_ to the I cells shifts the network dynamics from PING to ING in E-I spiking network simulations. **(A)** Raster plot for PING shows that both E and I populations are rhythmic, with I cells tightly synchronized and E cells loosely synchronized. The episode of I cell firing terminates the E-cell firing, setting the silent window of the rhythm. Each dot represents a spike of an E cell (red, cell index from 1 to 800) or an I cell (blue, cell index from 801 to 1000). The E cells are sorted by *I*_E_ in descending order. Inputs: uniform distribution on [0.38, 0.42] μA/cm^2^ for E cells; heterogeneous subthreshold drive to I cells, uniform distribution on [−1.05, −0.95] μA/cm^2^. **(B)** Raster plot for ING shows that the E cells are asynchronous, and the I cells are less synchronous than during PING in **A**. Input: same as **A** for E cells; heterogeneous suprathreshold drive to I cells, uniform distribution on [0.95, 1.05] μA/cm^2^. **(C)** Rhythmic activities in the synaptic fields during PING: Time courses of cell-averaged synaptic conductance *g*_syn_ (top) and synaptic currents *I*_syn_ (bottom) within (solid) or between (dotted) E and I populations for the raster plots in **A**. The amplitudes of the I-to-I synaptic conductance and synaptic current are dominant due to the high synchrony level of the I cells. E-to-I, I-to-E and E-to-E synaptic conductance and current have relatively small oscillation amplitudes. **(D)** As with **C** but for the raster plot in **B**. During ING, the oscillations in I-to-I synaptic conductance (current) are still most prominent among the coupling feature time courses but with a smaller amplitude than in PING, reflecting less synchrony of the I cells. The E-to-E synaptic field is almost constant over time due to near asynchrony among E cells. The I-to-E synaptic conductance (current) is too flat to recruit the E cells into the rhythm. **(E)** Top: Time courses of the instantaneous firing rate *R* (solid, window size = 5 ms), mean synaptic gating *s* (dashed) and mean membrane potential *V* (dotted) for the E population during PING. The dashed horizontal black line marks the asynchronous E cell rate (139 Hz) without I cells, indicating that this is the target rate for E cells during PING. Bottom: Similar with the top panel but for the I population. The firing rate *r* and the mean membrane potential *v* are normalized (See Methods 4.3 for details about normalization). The time courses illustrate the recurrent activation of the variables in the order of *r*, *s* and *v* for both E and I populations. The time course of <spk/cycle> shows the average number of spikes per cycle of the cells in corresponding color. **(F)** Same with **E** but for ING. The mean instantaneous firing rate for E cells modulates over a relatively small range, indicating that the E cells are nearly asynchronous and fire at near constant rate. The oscillation amplitude reflects the degree of synchrony, matching with the observation in **A** and **B** that both E and I cells are more synchronized in PING than ING.

In the rhythmic PING state (Fig 1A), the I cells receive subthreshold external input and therefore can only be recruited by the E cells. Their firing shuts down the E cells as well as their own firing, hence their firing episode is brief. The recovery of E cells has two phases: a gradual phase (during the first 15 ms after an I-cell episode) of slowly increasing E-cell firing from near silence as synaptic inhibition wears off followed by a rapid phase (5 ms preceding the next I-cell episode) where E cell firing accelerates because of recurrent excitation. This autocatalytic rapid rise in E-cell activity recruits I cells and gives rise to another I-cell episode. The reciprocal interaction between the E-I cells leads to the steady alternating PING-pattern of network activity: increasing activity of E cells that recruits nearly synchronous firing of I cells. The two-phase aspect of E-cell recovery can be understood by examining the dynamics of emergence and termination of PING during a duration of I-cell drive that suppresses I cells (S1 Fig).

On the other hand, ING arises when the I cells are sufficiently driven by suprathreshold input (Fig 1B) and the I cells show coordinated cyclic firing without needing temporally patterned synaptic input from the E cells. The nearly asynchronous E-cell firing provides an effectively constant synaptic field to the I cells. Since *I*_syn,IE_ is very small compared to *I*_syn,II_, the impact of the E cells on the ING firing pattern is not strong except for slightly increased oscillation frequency. The modest *I*_syn,IE_ can be viewed as contributing to the steady input drive *I*_I_ as if in an isolated I-I network. During ING, as compared to the preceding PING case, the I cells receive more total drive and they have higher mean cellular frequency (50.37 ± 0.79 Hz in ING versus 42.1 ± 0.83 Hz in PING). The mean cellular frequency of E cells is comparable in PING (53.67 ± 2.29 Hz) and ING (54.26 ± 3.22 Hz), which is much less than their asynchronous firing rate (139 Hz) in an isolated E-E network. The network oscillation frequency during ING (85.23 ± 5.12 Hz) is nearly twice that during PING (46.17 ± 1.94 Hz). The I to E inhibition and high ING frequency allow little chance for the E cells to form a significant rhythmic synaptic field since E cells cannot return to their asynchronous rate before the next I episode. Notably, the mean cellular frequency of I cells is almost half of the network frequency in ING, suggesting that many I cells skip cycles; but in PING, these two frequencies are close, suggesting that most I cells spike in each cycle.

Examination of the cell-averaged synaptic gating variables *g*_syn_ (Fig 1C top and 1D top) and currents *I*_syn_ (Fig 1C bottom and 1D bottom) provides further understanding of the factors that determine the frequency of PING and ING. During PING (Fig 1C), the cell-averaged time courses are clearly rhythmic. Most prominent are the highly synchronized I-cell firing episodes that generate a strong synchronized inhibitory synaptic output, *g*_syn,II_ and *g*_syn,EI_ with sharp onset and exponential decay *τ*_syn,I_. These conductance transients lead to currents *I*_syn,II_ and *I*_syn,EI_ (dashed blue, from I to E) that shut down I-cell and E-cell activity. Recovery of the E cells is set by the slower decay time of *g*_syn,EI_ and by the gradual re-establishing of asynchronous E-cell firing and *g*_syn,IE_ (S1 Fig, bottom panel, dashed red just after 140 ms, from E to I). In contrast, the oscillation in the synaptic fields *g*_syn,EE_ (solid red) generated by the E cells is weak due to the low level of synchrony amongst E cells. Notice that even though the rise of *g*_syn,IE_ is smooth, there is a sudden sink during the activation of *I*_syn,IE_ (dashed red). When the membrane potential of a postsynaptic I cell rises toward the reversal potential of an excitatory synapse, *I*_syn,IE_ is transiently reduced.

The amplitudes of *g*_syn_ and *I*_syn_ during ING (Fig 1D) are smaller than those during PING due to the reduced level of synchrony in both populations. Since the I cells are still moderately synchronized, prominent oscillations are seen in *g*_syn,II_ and *I*_syn,II_ (blue solid). In contrast, *g*_syn,EE_ and *I*_syn,EE_ (red solid) are almost constant over time, reflecting that the E cells are almost asynchronous. Although *g*_syn,EI_ shows noticeable modulation, the effect on *I*_syn,EI_ is lessened since the driving force for inhibition is small and, indeed, if we replace *g*_syn,EI_(*t*) with its time average the I cells still display a very similar ING pattern (S2 Fig).

**Fig 2.**
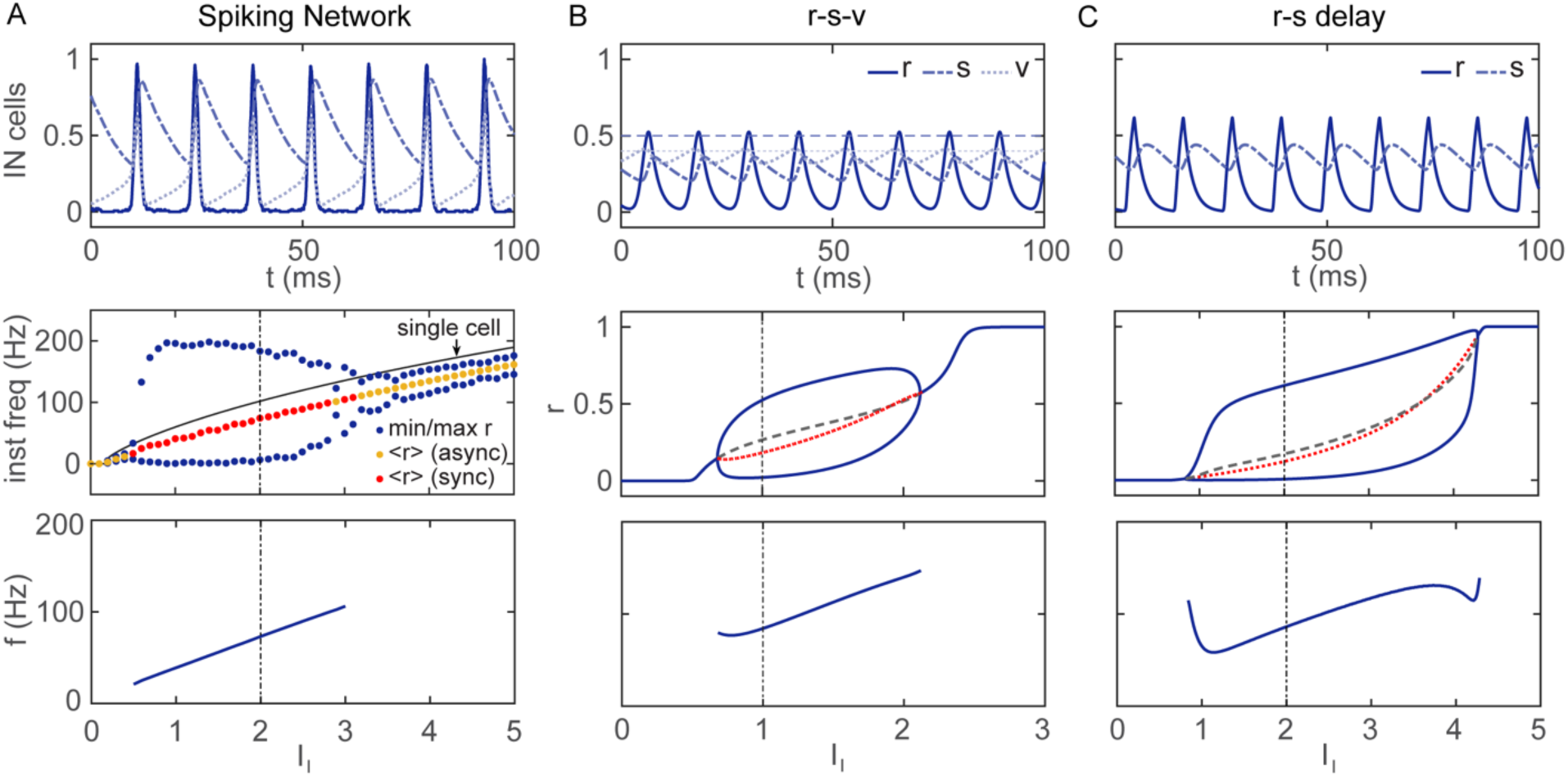
Firing rate models capture features of the inhibitory spiking network. **(A)** Spiking network consisting of 200 I cells. The single neuron model is from Wang & Buzsáki [27]. See Methods 4.2 for details about network set-up. Top: The sample normalized time courses of instantaneous firing rate (*r*, solid), mean synaptic gating variable (*s*, dashed) and mean membrane potential (*v*, dotted) show the activation of *v* is followed by *r* and then *s*. Here, the heterogeneous steady drive has mean *I*_I_ = 2 μA/cm^2^ (black vertical dashed line, bottom two panels) and standard deviation of 0.02 μA/cm^2^. See Methods 4.3 for the details in calculating the mean-field variables from spiking network simulation. Middle: Instantaneous firing rate versus *I*_I_ with rate calculated as # spikes/(# cells · window size) with window size = 5 ms; also plotted are maximum and minimum rate (blue dots), the mean rate (red dots); yellow dots correspond to no oscillation. Moderate input drive *I*_I_ is needed for ING rhythm generation. Bottom: The network oscillation frequency increases monotonically with *I*_I_. **(B)** The three-variable *r-s-v* firing rate model. See Methods 1.1 for parameter values. Top: The sample time courses for firing rate (*r*, solid), synaptic gating variable (*s*, dashed) and mean membrane potential (*v*, dotted) with *I*_I_ = 1 reproduce the activation order of *v*, *r* and *s* in the spiking network model shown in **A**. Horizontal dashed line and dotted line indicate the value of *θ_q_* and *θ_f_* respectively. Middle: The one-parameter bifurcation diagram, i.e., maximum (upper solid blue), minimum (lower solid blue) and steady states (solid blue for stable, dashed grey for unstable) of firing rate *r* versus *I*_I_, has two supercritical Hopf bifurcations, representing the onset and offset of oscillations. For comparison with the spiking network, the temporal mean of firing rate is shown (dotted red line). Bottom: The oscillation frequency versus *I*_I_ is almost monotonically increasing, similar with the spiking network model in **A**. **(C)** The two-variable *r-s* delay rate model with delay time constant *δ* = 1ms. See Methods 1.2 for model equations and parameter values. Panels as in **B** except that *I*_I_ = 2 for the time course (top). The *r-s* delay model shows minor differences with the *r-s-v* model: larger oscillation amplitude in *r* and nonmonotonic frequency dependence on *I*_I_.

In this example spiking network, PING and ING are bistable for some input drive *I*_I_ near the single I cell firing threshold (S3 Fig). When the network is in ING, the combined input *I*_I_ and nearly constant *I*_syn,IE_ to the I cells is always above the threshold. Since the E cells are suppressed and firing slowly (50 Hz) comparing to the asynchronous rate of an isolated E network (139 Hz), they are unable to synchronize and generate rhythm, the network remains in ING. However, when the network is in PING, even with the decay of *I*_syn,II_ to near zero, the subthreshold *I*_I_ means that I cells do not fire on their own and must wait long enough for *I*_syn,IE_ to rebuild and then activate synchronously the I cells. Somewhere between sub and suprathreshold *I*_I_ is a sweet spot for our network example so that both PING and ING can co-exist.

**Fig 3.**
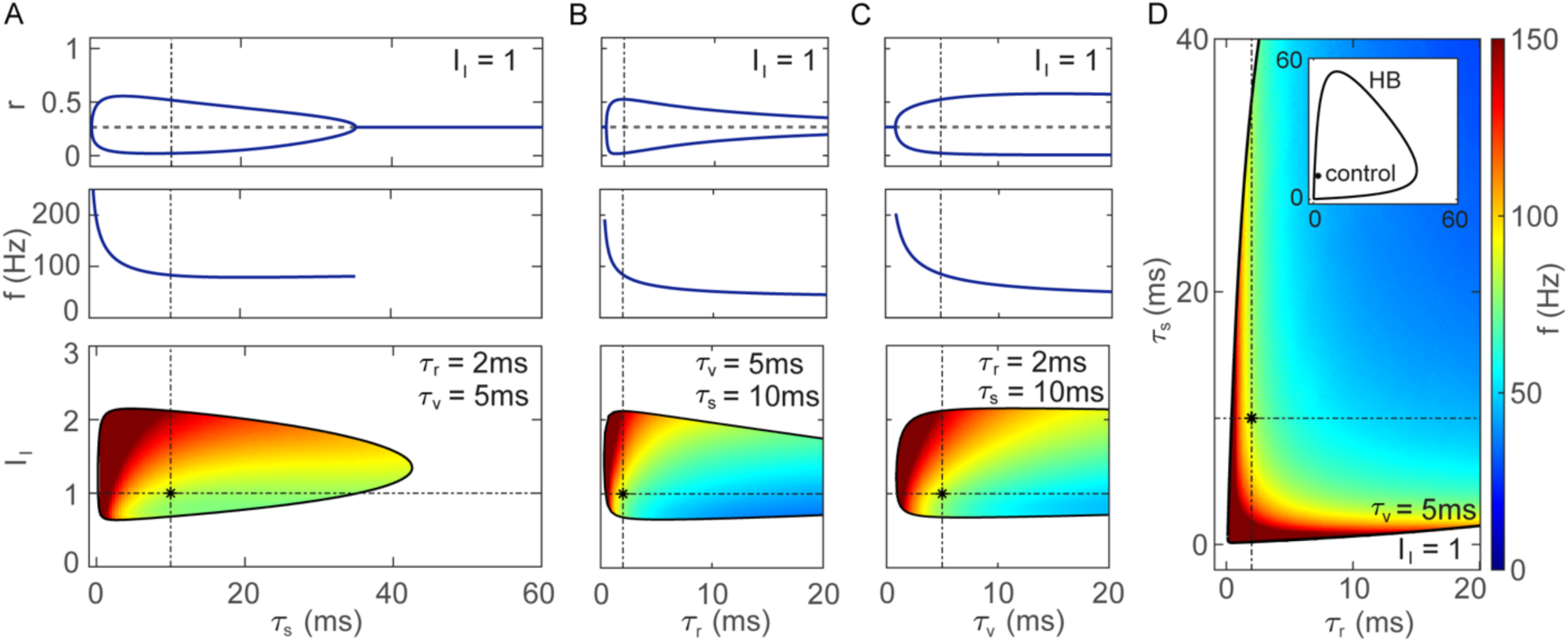
Dependence of the ING oscillation on the time constants. **(A)** Top: One-parameter bifurcation diagram, the firing rate *r* versus the time constant *τ_s_*, with two Hopf bifurcations for the onset and offset of oscillations indicating that *τ_s_* cannot be too small (not too fast) or too large (not too slow) for oscillation. Middle: The oscillation frequency decreases monotonically with *τ_s_* due to slower *s_I_*-dynamics and longer period. The frequency gradually plateaus as *τ_s_* gets large. Bottom: Two-parameter bifurcation diagram for *τ_s_* and *I*_I_ shows the oscillation regime delimited by Hopf bifurcations (black curve) and heat map for the oscillation frequency. The frequency increases monotonically as *τ_s_* decreases and *I*_I_ increases. For *I*_I_ too large or too small the steady state is stable. Control parameter values: black dashes and asterisk. **(B)** Same layout as in **A** but for the time constant *τ_r_* and different horizontal axis scaling. The Hopf bifurcation for oscillation offset at high *τ_r_* is omitted for better resolution in region of interest. The dependence on *τ_r_* has similar features as for *τ_s_*; *τ_r_* can neither be too large or small for oscillation. Notice that the oscillation amplitude shrinks rapidly as *τ_r_* increases, indicating weak modulation of firing rate (weak synchrony) unless *τ_r_* is relatively small. The network frequency decreases significantly as *τ_r_* increases and plateaus beyond a few ms. **(C)** Same layout as in **B** but for the time constant *τ_v_*. The dependence of *τ_v_* has similar features with *τ_r_*, except that high oscillation amplitude persists for a larger range of *τ_v_* and the plateau of network frequency is not as abrupt. **(D)** Two-parameter bifurcation diagram for *τ_r_* and *τ_s_* with *τ_v_* fixed at 5 ms shows the symmetry in the effects of the time constants. As either *τ_r_* or *τ_s_* increases, the frequency decreases. There is a trade-off between *τ_r_* and *τ_s_* in maintaining a constant oscillation frequency (same color). The inset shows a triangular-shaped Hopf boundary in the full parameter space with *τ_r_* and *τ_s_* sufficiently large (*τ_r_* and *τ_s_* in ms for the horizontal and vertical axis respectively).

Although a spiking network describes single cell activities and captures detailed information, it does not lend itself to analysis and it is computationally expensive especially for large-scale modeling [41]. In order to study the dependence of network dynamics on parameters efficiently, we seek a firing rate representation of PING and ING and to account for the PING to ING transition. To motivate the firing rate model that we develop below, we plot here mean-field-like averaged variables from the simulations of the spiking model: the instantaneous firing rate *R*_E_*, r*_I_, cell-averaged membrane potential *V*_E_*, v*_I_ and synaptic gating variable *s*_E_, *s*_I_ for E (top, with unit) and I (bottom, normalized) cells in the example PING (Fig 1E) and ING (Fig 1F) spiking network simulations (Fig 1A, 1B respectively). The amplitude of the firing rate oscillation reflects the degree of synchrony. Consistent with the raster plots, both the E cells and I cells are more synchronized in PING than in ING. Also, the nearly constant E-cell firing rate in ING is relatively lower than the I-cell rate. During ING, the activation of membrane potential *v*_I_ triggers the rise of firing rate *r*_I_, that subsequently activates the synaptic gating *s*_I_ that drives the inhibitory synaptic current to decrease membrane potential *v*_I_. Consequently, the firing rate *r*_I_ drops and then the synaptic gating *s*_I_ deactivates. The loop formed by membrane potential *v*_I_, firing rate *r*_I_ and synaptic gating *s*_I_ supports the repetitive generation of the oscillation.

The classical Wilson-Cowan (WC) model of E-I populations with additional dynamical variables for synaptic activation can generate PING oscillations [32,33], yet it is unable to capture ING dynamics. Indeed, from linear stability analysis it follows that the two-variable *r-s* WC model for a single inhibitory population always has one stable fixed point and therefore cannot generate an oscillation (see Methods 2 for the proof; see also [34]). So, one major challenge in developing a rate model that can account for both PING and ING is to have an ING-capable model for an inhibitory-only population. In the next section, we offer alternative formulations by including an auxiliary variable to an *r-s* rate model: either *v* as a mean-voltage variable or *u* as pre-synaptic variable that transforms spike rate *r* to synaptic activation *s*.

### 2 An *r-s-v* firing rate model for ING that reproduces responses of a spiking inhibitory network model

The standard *r-s* Wilson-Cowan model fails to capture ING dynamics because it lacks a delay mechanism and has no regenerative process; the firing rate directly activates *s* which immediately feeds back to suppress *r*. One resolution is to introduce an explicit delay for activating *s* as in the *r-s* delay model as shown in [32]. However, the explicit delay presents challenges for analytic treatment, so we propose involvement of an implicit delay. In the spiking network, explicit delay is not required for generating ING (although a delay is incorporated in several other models for ING, and found to render the behavior more robust, see e.g., [42]). The subthreshold build-up of single neuron membrane potential contributes to the delay activation of the synaptic gating and therefore is a candidate delay mechanism for a firing rate model.

Motivated by the spiking network simulation in Fig 1F, we introduced an additional dynamic variable into the traditional Wilson-Cowan model that describes the temporal integration of mean membrane potential *v* over cells. The model equations are given as follows.

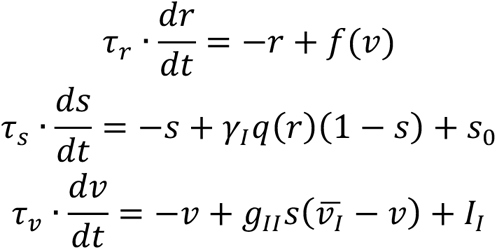

where the input-output function *f*(*v*) is a sigmoid function

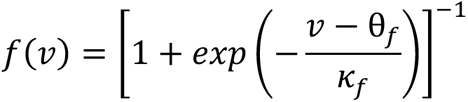

*q*(*r*) describes the dependence of the synaptic activation rate on the presynaptic firing rate by

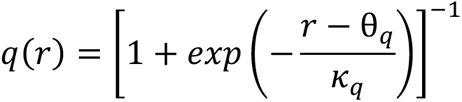

In the *r-s-v* model, the additional *v*-variable can represent the subthreshold integration of inputs and acts as an effective delay in the negative feedback loop between firing rate *r* and synaptic gating of inhibition *s*.

The three-variable *r-s-v* model for an I population captures many dynamic properties of ING in our simulations of an I-I network of Wang-Buzsáki cells [27], as developed in their systematic study of conditions for coherent gamma oscillation in an I-I spiking network. We find striking similarities in comparing the time courses of averaged firing rate, membrane potential and synaptic gating variable from the spiking network simulation with the analogous variables *r*, *v*, *s* respectively in firing rate model. In the isolated I-I network (Fig 2A top), the time courses for I cells are similar with those in the spiking E-I network during ING (Fig 1F). The temporal sequencing of *v*, then *r* and then *s* is successfully reproduced in the firing rate model (Fig 2B top). The amplitude differences in the transients between the rate and spiking models are not concerning since there is no uniform rule for normalization in these two models and the amplitude of instantaneous firing rate of the spiking network depends sensitively on the window size. Moreover, the level of synchrony of the spiking network depends on factors such as heterogeneity in input drive and connectivity, while the firing rate model, by its nature, assumes the heterogeneity without a quantification. However, one difference is worth noticing between the models. In the spiking network model, the firing rate is pulsatile and drops to near zero between spike clusters, matching with the fact that the I-cells are silent. This is not the case in the *r-s-v* firing rate model. There, the firing rate is driven by *f*(*v*). After a firing event and before *v* gets small enough such that *f*(*v*) ≈ 0, allowing *r* to decay to zero and flatten out, the inhibitory activation *s* decays enough so that the subthreshold *v* starts increasing, able to activate the next cycle. We also compare the time course of *r-s-v* model with the *r-s* delay model (Fig 2C) and observe the similarity except that *r* flattens a bit between I-firing events. With long explicit delay (not shown), the *r-s* delay model can achieve the pulsatile firing rate as in the spiking network.

We next compared the ING oscillation’s dependence on the input *I*_I_ in these three models. Here, we focus on the temporal maximum, minimum and mean of the variable *r* (Fig 2 middle) and the network frequency (Fig 2 bottom) as the representative features. The synaptic gating variable *s* and mean membrane potential *v* have similar characteristics (S4 Fig). For the spiking model, when *I*_I_ is small (< 0.68), the network shows sparse asynchronous firing (Fig 2A middle). With moderate *I*_I_, the cells are above threshold and the recurrent inhibition restricts the firing to a brief window when *I*_I_ + *I*_syn,II_ is at a minimum – the network is in ING oscillation. As *I*_I_ increases, the intrinsic frequency of each cell also increases, leading to faster network rhythms (Fig 2A bottom). When the input drive *I*_I_ is large enough (> 2.12), it dominates over the synaptic currents. Temporal organization is compromised; the network is in asynchrony and cells fire according to their individual input *I*_I_ and the mean synaptic field. The firing rate model reproduced the observations from the spiking network. For the rate model, we can identify the onset and offset of ING oscillation as due to two supercritical Hopf bifurcations (Fig 2B middle). These correspond to weak modulation of firing in the spiking network (not shown, more in Discussion). The ING oscillation frequency increases with input drive as for the spiking model (Fig 2B bottom). The *r-s* delay model has similar characteristics as the *r-s-v* model in terms of the firing rate (Fig 2C middle), but the network frequency has a nonmonotonic dependence on *I*_I_ (Fig 2C bottom).

### 3 Dependence of ING oscillation on parameters in the *r-s-v* firing rate model

To demonstrate the robustness of the *r-s-v* firing rate model, we explored the dependence of the ING oscillation on the time constants (Fig 3) and other synaptic parameters such as the maximal conductance and the reversal potential (Fig 4).

**Fig 4.**
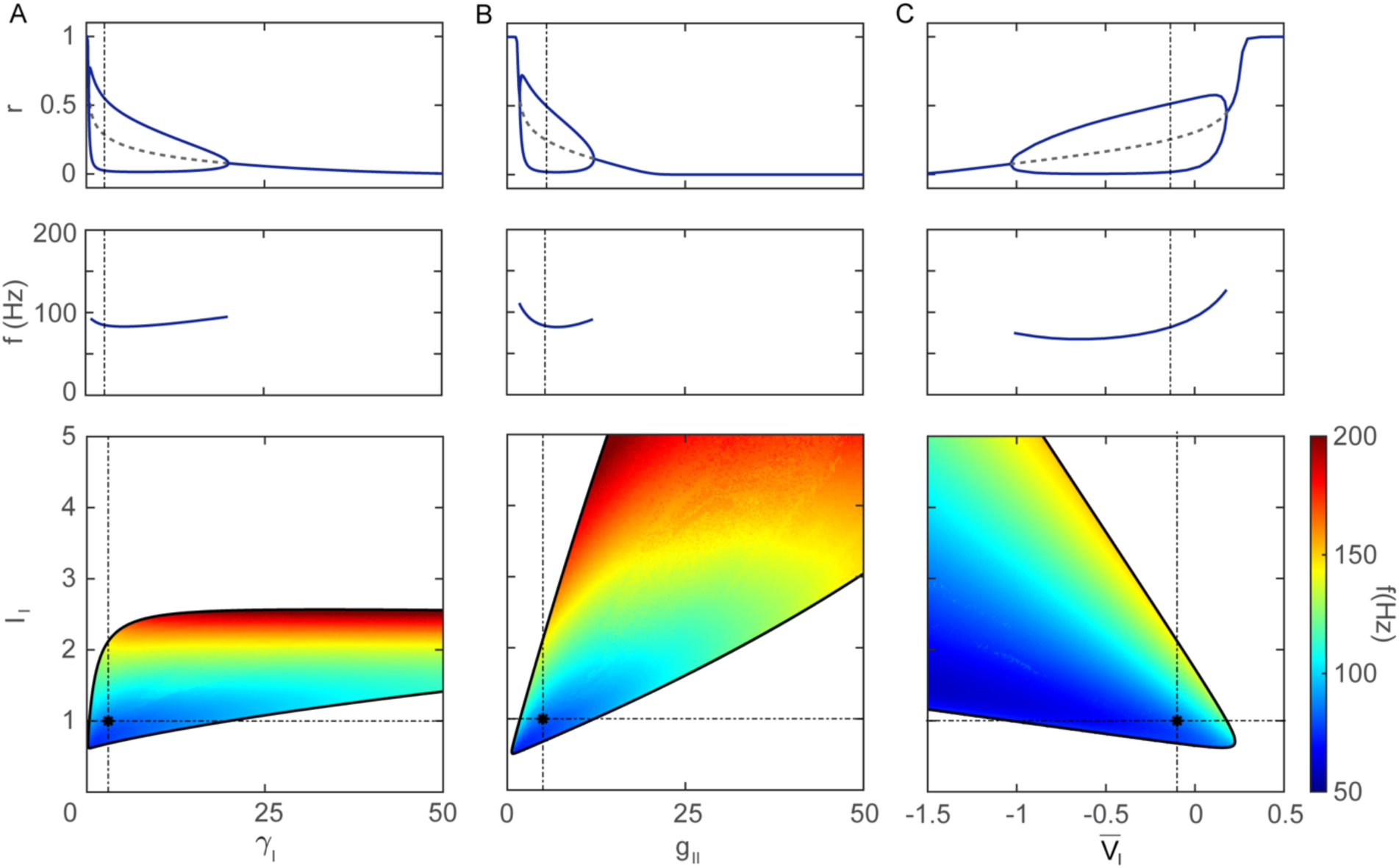
Dependence of the ING oscillation on synaptic properties. **(A)** Effect of the synaptic activation rate *γ*_I_. Notice that the synaptic rise-time constant *τ_s_* /*γ*_I_ scales inversely with *γ*_I_. Panel layouts and notations are as in Fig 3. Top: Two Hopf bifurcations show that a moderate delay in synaptic activation is needed for oscillation. Middle: The oscillation frequency depends only weakly on the synaptic rise-time constant. As *γ*_I_ increases (rise-time constant *τ_s_* /*γ*_I_ decreases) tending toward zero it has little effect on the oscillation period. Bottom: The *I*_I_ -range for oscillation shrinks a bit then stabilizes as *γ*_I_ increases. For a fixed *I*_I_, the frequency varies little with *γ*_I_; the color bands are horizontal, as is the plot in the middle panel. **(B)** Effect of the recurrent inhibitory conductance *g*_II_. Top: Sufficient inhibitory synaptic conductance is required for stable synchronization of the network. However, too much recurrent inhibition will destroy the synchrony by inhibiting the I population. Middle: The oscillation frequency first decreases and then increases with *g*_II_. Bottom: Stronger applied current can compensate the increased recurrent inhibition, while leading to faster oscillation. **(C)** Effect of the inhibitory reversal potential 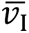. Top: Hyperpolarizing inhibition (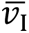 < *v*_rest_ = 0) promotes oscillation rather than shunting inhibition (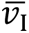 ≥ *v*_rest_ = 0). Although there are two Hopf bifurcations in the diagram, we consider a very negative 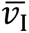 value as beyond the range of biophysical interest. Middle: The oscillation frequency increases with 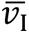 over the range of interest. Bottom: As *I*_I_ increases, the Hopf bifurcation for the offset of the oscillation is shifted to the left, indicating that hyperpolarizing inhibition is more favored than shunting inhibition.

The bifurcation diagrams for the time constants *τ_s_*, *τ_r_* and *τ_v_* (Fig 3A, 3B, 3C respectively) show that for the existence of ING no time constant can be too slow nor too fast (set to zero). In other words, the model cannot be reduced to a two-variable model by eliminating one variable by treating is as extremely slow or extremely fast. For each time constant, the onset and offset of oscillations occur by supercritical Hopf bifurcations. Between the Hopf bifurcations, the oscillation frequency decreases and plateaus as each time constant increases. The gradation of frequency is broad over *τ_s_* as we have come to expect for oscillations of inhibition-only networks (Fig 3A): the synaptic decay rate robustly sets the oscillation frequency. The oscillation amplitude decreases dramatically with *τ_r_*, yet not with *τ_s_* and *τ_v_* (Fig 3B), indicating that ING will be weak if recruitment of the I population is too slow, consistent with our intuition. The control of ING frequency by input drive is more labile/robust over the allowable range of *τ_s_*, than for *τ_r_* and *τ_v_* (Figs 3A, 3B, 3C bottom). The two-parameter bifurcation diagram of *τ_r_* and *τ_s_* (Fig 3D) show the complementary role of the time constants. To attain a fixed oscillation frequency, there is a trade-off between *τ_r_* and *τ_s_* – increasing either would require a compensatory decrease of the other. This diagram would not change shape with a different value of *τ_v_*; only the oscillation frequency (a simple scaling) would change but the existence and amplitude of the oscillation would not be affected.

The numerical observations can be validated by linear stability analysis of the three-variable system (see Methods 3). We first showed that an exchange of the fixed point’s stability can change only by Hopf bifurcation. Therefore, to ensure the system has a stable limit cycle solution for an ING oscillation, Hopf bifurcation must exist. Then by the trace condition for a Hopf bifurcation, we derived a necessary condition for oscillation given by

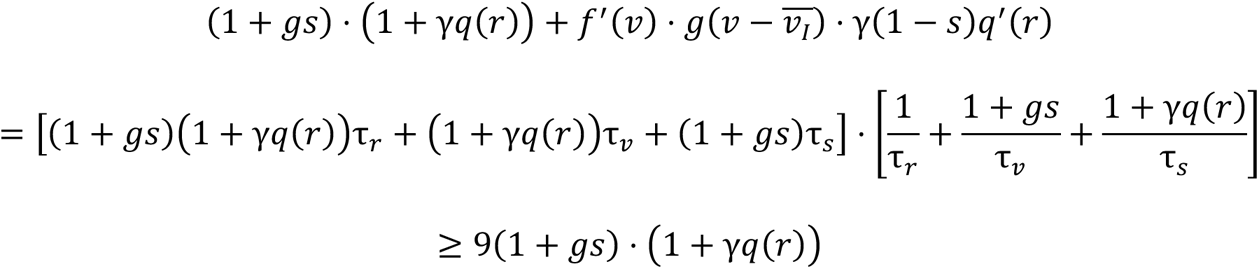

The equality is attained when the effective time constant of the three variables are the same, i.e.

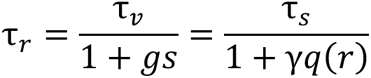

This confirms that none of the time constants can be zero or infinity, suggesting the existence of two Hopf bifurcations with respect to each time constant.

We further examined how other synaptic dynamics related factors influence the existence properties of ING in the rate model, in particular the effect of the synaptic activation time constant *τ_s_* /*γ*_I_, the maximal inhibitory synaptic conductance *g*_II_ and inhibitory reversal potential 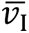_+_ on ING in the firing rate model (Fig 4).

Firstly, the synaptic rise-time constant *τ_s_* /*γ*_I_ has a stronger effect on the ING amplitude than on the frequency; the frequency is determined more by the slower decay-time constant *τ_s_*. For moderate input drive *I*_I_ the oscillation amplitude decays with *γ*_I_ (Fig 4A top, middle). If the rise is too fast the rapid activation of inhibition shuts down firing and precludes ING. For stronger input the allowable range for *γ*_I_ increases but still has little impact on frequency (Fig 4A bottom).

Secondly, the maximal inhibitory synaptic conductance *g*_II_ also needs to be moderate (Fig 4B top). If recurrent inhibition is too weak, there is no oscillation, interpretable as failure to synchronize in the I population, while too strong recurrent inhibition suppresses the activity of the I population, i.e. very low firing rate. These effects match with the observations in spiking network model [27]. The oscillation frequency has a non-monotonic dependence of *g*_II_, where the minimal frequency is attained with a moderate synaptic conductance (Fig 4B middle). The loss of oscillation at high *g*_II_ can be compensated by increased *I*_I_ (Fig 4B bottom). As *g*_II_ and *I*_I_ increase together, the amplitudes of the input drive and the synaptic current both grow but remain comparable, leading to an increase in network frequency.

Thirdly, hyperpolarizing inhibition rather than shunting inhibition benefits an ING oscillation. In the rate model, the mean membrane voltage variable *v* is normalized such that the rest potential *v*_rest_ is zero and the excitatory reversal potential satisfies 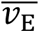 = 1. Assuming, in unscaled variables, *V*_rest_ = −65 mV and 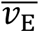 = 0 mV, the control inhibitory reversal potential is given by 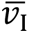 = −0.1 corresponds to 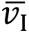 = −71.5 mV. In the one-parameter bifurcation diagram of the inhibitory reversal potential 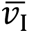 (Fig 4C top), the Hopf bifurcation with large 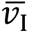 value shows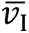 cannot be too high above the rest potential, *v*_rest_ = 0, for oscillation. The Hopf bifurcation with small 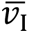 value is far below a biological realistic value and not of interest. In the biologically realistic regime (say 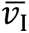 > −0.5), the oscillation frequency increases monotonically with the inhibitory reversal potential (Fig 4C middle). With fixed 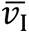 (Fig 4C bottom, vertical slices), the *I*_I_ range for oscillation shrinks and disappears as 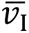 gets to around 0.2. As *I*_I_ increases, the Hopf bifurcation with large 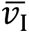 value decreases below 0 (Fig 4C bottom, horizontal slices), indicating only hyperpolarizing inhibition supports oscillation.

## 4 The *r-u-s* model for ING with implicit delay in synaptic activation

Here, we continue to demonstrate the importance of introducing a delay in an *r-s* rate model in order to generate gamma oscillations. In the previous section we described the *r-s-v* model, in which the mean voltage variable *v* acts as an implicit delay between the negative feedback from the synaptic gating *s* to firing rate *r*. The variables *r*, *s* and *v* form a loop to generate the oscillation recurrently in the I-I network (Fig 5D). Alternatively, we consider here an implicit delay by introducing an intermediate variable *u* between the presynaptic firing rate *r* and the activation of synaptic gating *s*. The variable *u* can be thought of as representing dispersion of arrival times of spikes onto target neurons or a preparatory process prior to synaptic activation or a combination of such dynamic processes. In our *r-u-s* model, *r* activates *u*, *u* activates *s* and *s* sends negative feedback to *r* to form a loop (Fig 5A). The effective delay by *u* works more like the explicit *r-s* delay model. The model equations are

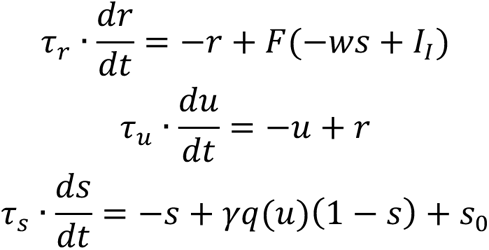

where *F* and *q* are both sigmoid functions as in the *r-s-v* model (see Methods 1.3 for parameter values).

**Fig 5.**
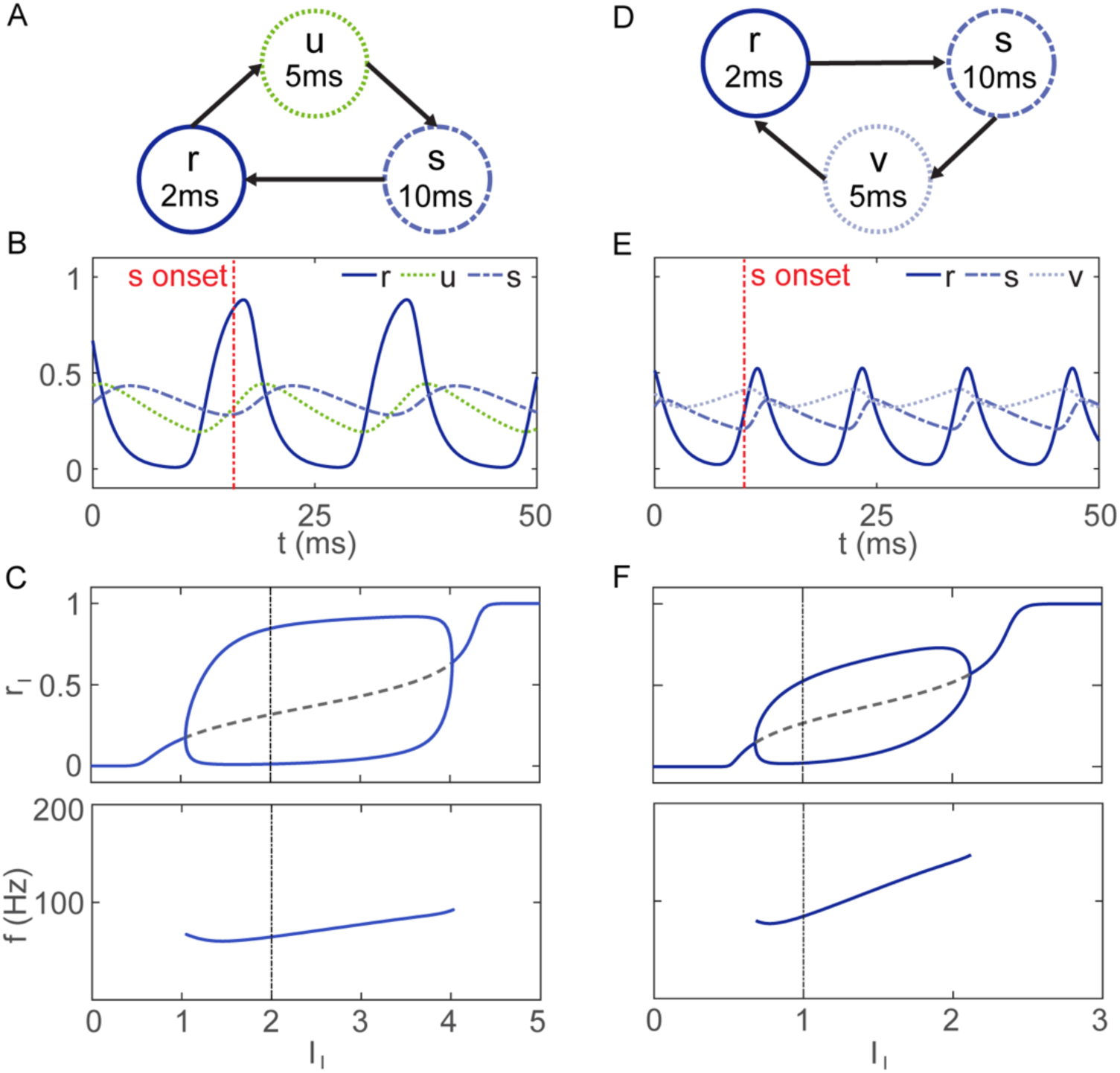
The *r-u-s* model and comparison with the *r-s-v* model. **(A)** Schematic for the *r-u-s* model shows that *u* acts as an effective delay from *r* to *s*. **(B)** The time course of the *r-u-s* model (*κ_F_* = 0.05 and *I*_I_ = 2) shows that the firing rate’s oscillation amplitude exceeds that in the *r-s-v* model (panel **E**) even though *κ_F_* is not so small. The synaptic activation does not begin rising until nearly the peak-time of firing rate due to the delay by *u* (compare the red vertical dashed lines in panels **B** and **E**). The firing rate stays near 0 for 5-10 ms before the next cycle begins. **(C)** The bifurcation diagram of firing rate and network frequency with respect to *I*_I_ of the *r-u-s* model shows a robust large amplitude and relatively low frequency oscillation in a wide *I*_I_ range compared to that for the *r-s-v* model (panel **F**). **(D)** Schematic for the *r-s-v* model shows that *v* acts as a delay from *s* to *r*. **(E)** The time course of the *r-s-v* model (*κ_f_* = 0.01 and *I*_I_ = 1) shows the synaptic activation happens quickly after the firing rate *r* crosses the threshold. **(F)** Repeated here from Fig 2B (middle and bottom panels) for ease of comparison.

Indeed, the example time course of the *r-u-s* model (Fig 5A) shows activation of variables in the order of *r*, *u* and *s*. One major difference between the *r-u-s* model and the *r-s-v* model is the silent phase between cycles. In the *r-u-s* model, the synaptic activation *s* does not respond to the rise of firing rate *r* directly, creating a time window for −*ws* + *I*_I_ to remain far below threshold and therefore *r* stays near 0. From the time courses, *s* increases quickly as *r* crosses the threshold in the *r-s-v* model (Fig 5B) and *s* peaks before *r* falls to one-half its maximum value; while in the *r-u-s* model, *s* begins to increase later, when *r* nearly reaches its peak, and *s* doesn’t peak until *r* is near to its lowest level. Moreover, in the *r-u-s* model, the oscillation amplitude is higher and consequently the frequency is lower. This is because the firing rate *r* has longer time to increase before the synaptic gating variable *s* is activated and begins to turn it off. The *r-u-s* model depends on the input drive *I*_I_ similarly to the *r-s-v* model, but with a larger amplitude for firing rate (Fig 5C) and with a larger dynamic range (*I*_I_ ∈[1.05, 4.03]) that spans lower gamma frequencies (*f* ∈[60.2, 92.4] Hz). A major parameter-dependent difference between the models is on the steepness of the input-output function as we describe below.

We find that for generation of ING oscillations the input/output functions and synaptic activation function of our rate models cannot be shallow. We describe the conditions about steepness, first, with numerical calculation and then obtain insight from an analytic treatment. Using the steepness (i.e., 1/(4*κ*), see Methods 3.3 for details) as a control parameter, we see a Hopf bifurcation that sets the maximal *κ_f_* (Fig 6A) or *κ_F_* (Fig 6C) for oscillation, indicating that the input-output function should be steep enough (*κ_f_* or *κ_F_* small enough). Notice that the allowable range of steepness for the *r-u-s* model (near 0.15) is almost 6 times that for the *r-s-v* model (near 0.025), indicating a significantly relaxed steepness constraint for the *r-u-s* model. In both models, the oscillation amplitude (peak firing frequency during an ING cycle) decreases as the input-output function’s steepness decreases (Fig 6A bottom for *r-s-v* model and Fig 6C bottom for *r-u-s* model). The reason is that a steep *f* or *F* leads to a faster change (large d*r*/d*t*) in the firing rate *r* and thus a larger oscillation amplitude. As a consequence, more time is needed for *s* to decrease enough so that the synaptic inhibition is small enough and the next cycle of spiking can occur. In terms of the change in the input drive *I*_I_, the frequency of the *r-u-s* model is more robust than that of the *r-s-v* model (Fig 6D versus Fig 6B). This is again due to the steepness difference in the input-output function. In the *r-s-v* model, *I*_I_ drives the membrane voltage *v*. As *I*_I_ increases, *v* passes through *θ_f_* from below and *r* increases quickly. While in the *r-u-s* model, *I*_I_ is part of the input argument to the less-steep function *F*, so the same change in *I*_I_ will lead to a smaller change in the magnitude of *r* and thus a smaller change in frequency.

**Fig 6.**
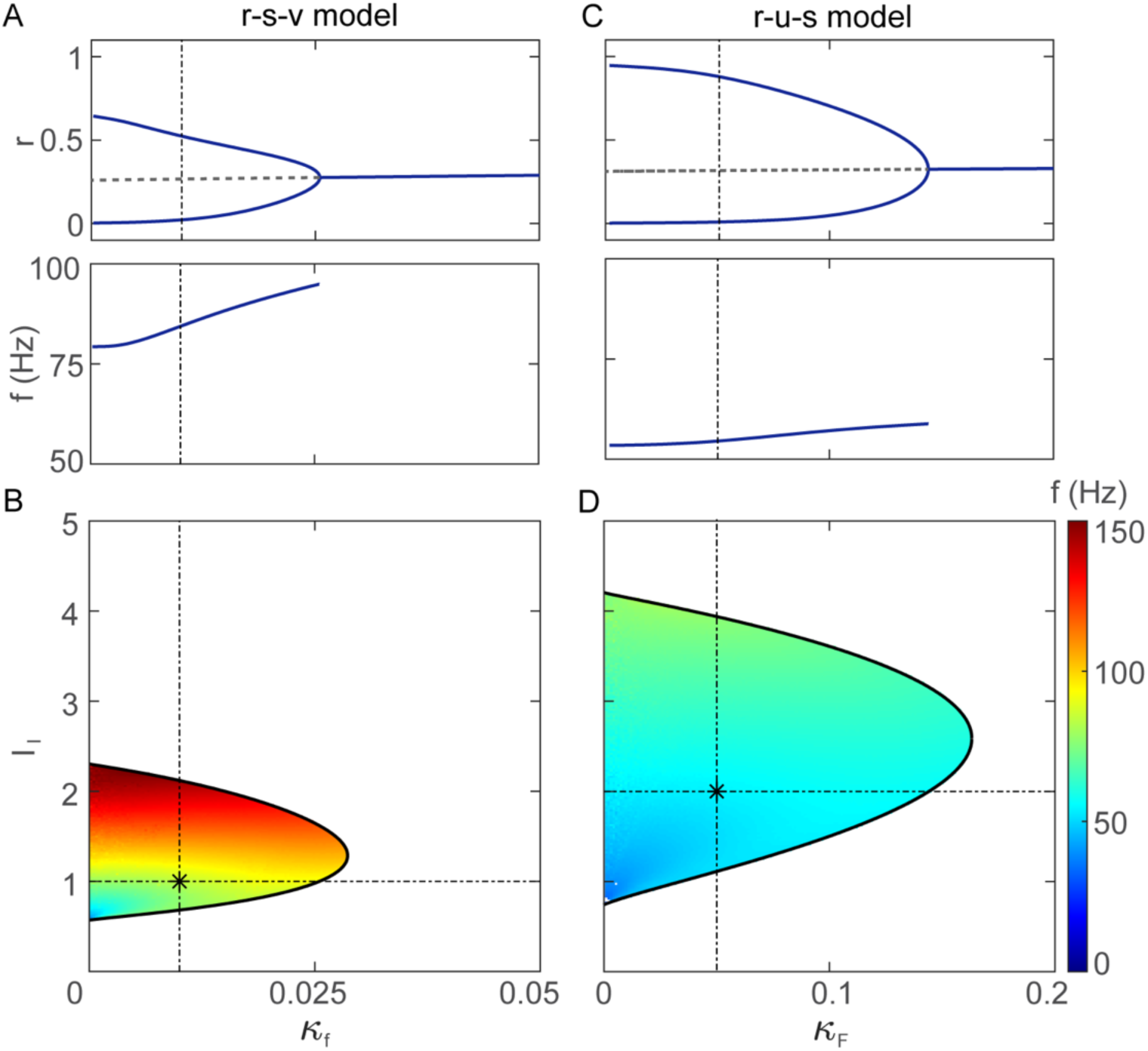
The effect on ING of input-output function steepness in the *r-s-v* and *r-u-s* models. **(A, B)** Bifurcation diagram of the *r-s-v* model with control parameter *κ_f_*, reciprocal of the of input-output function steepness. **(A)** Top: One-parameter bifurcation diagram, *r*_I_ versus *κ_f_*, shows that gamma oscillation terminates as a Hopf bifurcation. The steepness (1/4*κ_f_*) of the input-output function *f* scales inversely with *κ_f_*. Steep *f* corresponds to larger oscillation amplitude. Bottom: The oscillation frequency increases as *κ_f_* increases (*f* becomes less steep). **(B)** Two-parameter bifurcation diagram for *κ_f_* and *I*_I_ illustrates the restricted regime for oscillation where *κ_f_* values must be small. **(C, D)** Bifurcation diagram of *r-u-s* model with control parameter inverse steepness *κ_F_*. **(C)** Top: The oscillation regime of *κ_F_* is greatly extended in the *r-u-s* model from the *r-s-v* model. Bottom: Oscillation frequency in the *r-u-s* model is relatively low and has weak dependence on *κ_F_* compared to the *r-s-v* model. **(D)** The two-parameter bifurcation diagram illustrates an extended oscillation regime of the *r-u-s* model in the parameter space of *κ_F_* and *I*_I_.

Although the input-output function seems to be extremely steep for the *r-s-v* model, we show, by converting the dimensionless parameter values back to values with units, that the steepness feature is not biophysically implausible. As a sanity check, at the Hopf bifurcation, the slope of *f* can be approximated by 1/(4 × 0.025) = 10, so the steady state firing rate *r = f*(*v*) increases from 0 to 1 as *v* increases 0.1. Following the assumption that *V*_rest_ = −65 mV and 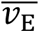 = 0 mV with units correspond to *v*_rest_ = 0 and 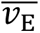 = 1 without units, a 0.1 increment in unitless *v* means a 6.5 mV increment in *V* with unit. Considering the heterogeneity in input drive and connectivity, it is reasonable that the firing thresholds of the cells lie in a range of 6.5 mV.

From the analytical perspective, we also derived for the *r-s-v* model a steepness condition of *f* for ING oscillation from the necessary condition of Hopf bifurcation (see Methods 3.2). Regardless of the time constants, we noticed that

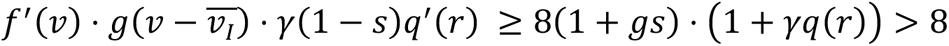

If *q*(*r*) is linear, as often assumed in such formulations *q*(*r*) = *r*, then we see from *s* > 0 that

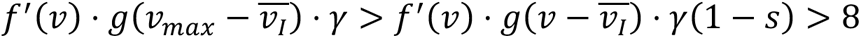

where *v*_max_ = max *v*(*t*) is finite. Therefore, *f* ′(*v*) is bounded from below, indicating that *f*(*v*) needs to be steep. The corresponding analysis applies similarly to *r-u-s* model (not shown). In the *r-s* delay model, the steepness constraint is considerably relaxed. It can be shown both numerically (S5 Fig) and analytically (S6 text) that as the explicit delay increases, less steepness in input-output function is required for oscillation.

## 5 A six-variable model for an E-I network and the PING-ING transition

Having developed the building block of ING in the I-I network, we extended the three-variable model for the I population to a six-variable model for the E-I populations to model both PING and ING. By repeating the I-equations and adding synaptic currents terms, the six-variable *r-s-v* model for E-I populations can be written as

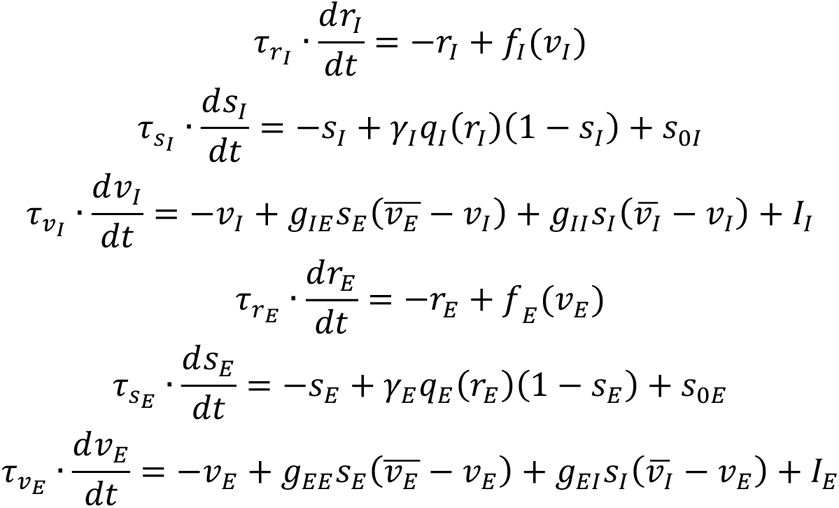

where the E and I populations have different thresholds and degrees of steepness for the functions *f* and *q* (see Methods 1.4). The six-variable system can be split into two subsystems, where the E and I subsystems are described by the last and first three equations respectively.

Each subsystem acts on the other only by the conductance-driven synaptic coupling, i.e., *s*_E_ drives *v*_I_ and *s*_I_ drives *v*_E_.

Without fine tuning the parameters, we can reproduce with the firing rate model (Fig 7) the transition from PING to ING by increasing the input drive *I*_I_ as seen in the spiking network (Fig 1). The example time courses for the E (top) and I (bottom) populations for PING (Fig 7A) and ING (Fig 7B) share similar features with the spiking network (Fig 1E and 1F). Firstly, the oscillation amplitudes of both E and I populations are larger in PING than ING. During PING, the firing rates of E and I populations are both oscillatory with the activation of E population earlier than the I population. While for ING, only the firing rate of the I population shows evident oscillation while the E population has low firing rate with almost no oscillation. The reduced oscillation amplitude of I population’s firing rate in ING reflects the lower degree of synchrony compared with that in PING. Secondly, the network oscillation frequency of ING is faster than that of PING. As we discussed for the spiking network, the oscillation period for PING involves two phases – the decay of synaptic inhibition and the gradual activation of the E population. That said, we find one notable difference between the rate and spiking models: the activation of the E population in the firing rate model takes longer than in the spiking network model.

**Fig 7.**
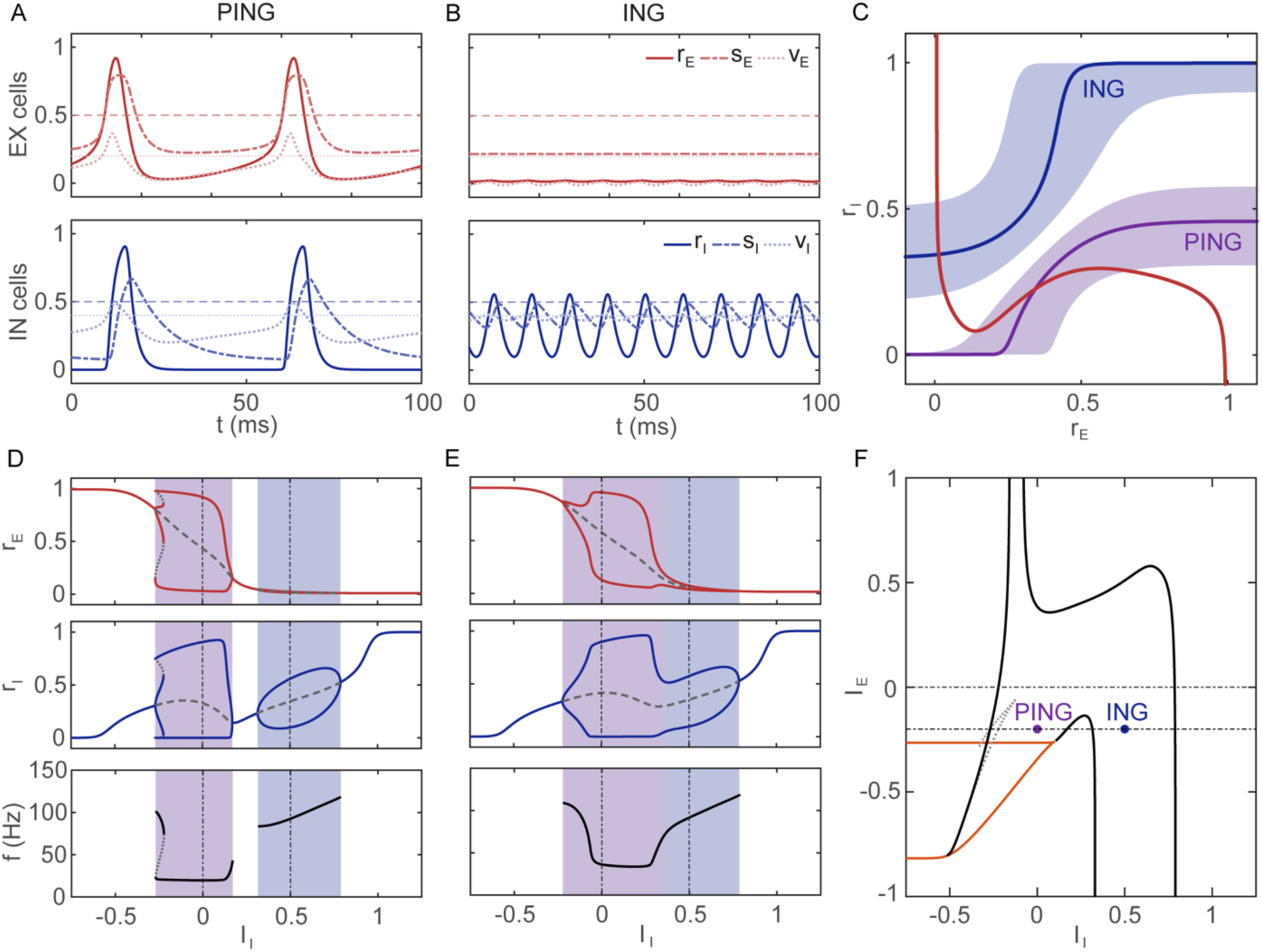
Six-variable *r-s-v* model for the E-I network. **(A)** Example time courses for PING (*I*_E_ = −0.2, *I*_I_ = 0). Top: Time courses for the E population (red) of normalized firing rate, *r* (solid), synaptic gating, *s* (dashed), and mean membrane potential, *v* (dotted), (*v* is normalized: 0 for rest potential and 1 for E reversal potential) show large amplitude oscillations. Bottom: Corresponding time courses for the I population (blue) also display large amplitude oscillations. The turn-on of excitatory firing rate *r*_E_ activates the excitatory synaptic gating *s*_E_ and depolarizes the I population. As *v*_I_ passes above threshold (thin dashed blue horizontal line), *r*_I_ rises and then *s*_I_ follows. The synaptic inhibition to *v*_E_ terminates the E-activity and *r*_E_ decreases. As *s*_I_ decays, *v*_E_ increases above threshold to begin the next cycle, leading to an oscillation of frequency 19.70 Hz that involves both E and I populations. **(B)** Same as in A but here, with stronger drive to I population, for ING (*I*_E_ = − 0.2, *I*_I_ = 0.5). Top: The E population is almost silent with almost no oscillation in firing rate. Bottom: The I population oscillates at 92.59 Hz with smaller amplitude (lower degree of synchrony) than during PING. **(C)** The projections of the *r*_E_ (thick red) and *r*_I_ (thick purple and blue) - SSRs in the *r*_E_ − *r*_I_ plane for the PING example (in **A**) and the ING example (in **B**) respectively show that PING operates on the middle branch of the *r*_E_ - SSR where both E and I populations are active; while ING operates on the left branch of the *r*_E_ - SSR, where E population is at very low (near zero) activity and I population is oscillatory. The purple and blue shades for PING and ING correspond to those in **D**. The boundaries of the shaded areas are the *r*_I_ - SSRs’ for *I*_I_’s at the Hopf bifurcations. **(D)** Bifurcation diagrams for *I*_I_ with *I*_E_ = −0.2 as in **A-C** has 4 Hopf bifurcations. As *I*_I_ increases, the network dynamics shift from PING (purple shade) to ING (blue shade) with separated regimes. Top: *r*_E_ has large oscillation amplitude in the PING regime and near zero oscillation amplitude in the ING regime. Middle: *r*_I_ has large oscillation amplitude in the PING regime and reduced oscillation amplitude in the ING regime. Bottom: The oscillation frequency is smaller in PING than ING. **(E)** Same bifurcation diagrams as in **D** but for *I*_E_ = 0 there are only two Hopf bifurcations. The PING-ING boundary as defined by the middle two Hopf bifurcations for low *I*_E_ has vanished upon their merging, but an effective transition can still be distinguished from the oscillation amplitude and frequency. **(F)** Two-parameter bifurcation diagram of *I*_E_ and *I*_I_ shows that the PING-ING transition by increasing *I*_I_ persists for a reasonable regime in the parameter space.

With these two examples for small and large *I*_I_, we next explored how the network dynamics varies over a range of input values, *I*_I_. The one-parameter bifurcation diagram for *I*_I_ shows four Hopf bifurcations (Fig 7D). The two Hopf bifurcations for smaller *I*_I_ (left) correspond to the onset and offset of PING, and the other two, for larger *I*_I_ (right), correspond to the onset and offset of ING. The oscillation amplitude and frequency of firing rates in PING vary little with the input drive *I*_I_ in comparison to ING. In the ING case, as we found in the three-variable *r-s-v* model, the oscillation frequency grows with *I*_I_ and the oscillation amplitude takes the maximal value when *I*_I_ is moderate. The oscillation amplitudes of synaptic gating variable *s*_I_ and the mean membrane potential *v*_I_ show notable differences in PING and ING as does the inhibitory firing rate *r*_I_ (S7 Fig).

To gain further understanding about the transition between PING and ING, we looked into the *r*_E_− *r*_I_ plane (Fig 7C). By setting all other variables to their steady state values as functions of *r*_E_ and *r*_I_, the *r*_E_ and *r*_I_ - steady state relationships (SSRs) can be projected into the *r*_E_ − *r*_I_ plane (see Methods 1.5 for details). The intersections of these SSRs correspond to steady states of the full system and the visualization has interpretive values although provides no information about stability. The shape of the SSRs has some basic and expected features as in nullclines of the standard two-variable *r*_E_ − *r*_I_ Wilson-Cowan model: the *r*_E_ - nullcline is cubic-shaped due to the regenerative effect of recurrent excitation; the *r*_I_ - nullcline increases monotonically with *r*_E_. With *I*_E_ fixed, the *r*_E_ - SSR (red) is also fixed. As *I*_I_ increases such that the network oscillation switches from PING to ING, the *r*_I_ - SSR is shifted towards the upper-left direction from the purple curve for PING to the blue curve for ING. The boundaries of the purple and blue shaded area are the *r*_I_ - SSRs for the *I*_I_ values at the four Hopf bifurcations in Fig 7D. It can be seen that the *r*_E_ and *r*_I_ - SSRs intersect on the middle branch of the *r*_E_ - SSR in the PING regime, while on the left branch of the *r*_E_ - SSR in the ING regime. Since the E population firing rate *r*_E_ is almost constant near 0 on the left branch of the *r*_E_ - SSR, the synaptic gating *s*_E_ is not activated and the mean membrane potential *v*_E_ is also near the rest value 0. In other words, when the I population receives large input drive *I*_I_, their activities are so strong that the E population is silenced by the synaptic inhibition. In this case, the I population works on its own to produce an ING rhythm without recruiting the E population.

For a range of *I*_E_ values, the PING and ING regimes are separated in the parameter space of *I*_I_ by Hopf bifurcations corresponding to their own onset and offset (as in Fig 7C and 7D). For somewhat larger *I*_E_, the PING and ING regimes are merged; at a critical value of *I*_E_, the middle two Hopf bifurcations collide and disappear (Fig 7E and 7F). Nevertheless, an effective transition for PING and ING dynamics is distinguishable based on the sudden change of the oscillation amplitudes of E and I firing rate. Again, the oscillation frequency is low at an almost constant value in the PING regime and increases monotonically to high values in the ING regime. The two-parameter bifurcation diagram for *I*_E_ and *I*_I_ validates the robustness of the PING-ING transition phenomenon in the parameter space (Fig 7F). For sufficiently low values of *I*_E_, the left trough of the *r*_E_ - SSR descends enough to intersect the *r*_I_ - SSR to create multiple steady states (saddle-node bifurcation) thereby disrupting oscillation mode of the PING regime; with extremely large *I*_E_, oscillations in the ING regime terminate by collision of the two Hopf bifurcations on the right. These dynamics for extreme values of *I*_E_, are not of primary interest here, so we only show some example one-parameter bifurcation diagrams for *I*_I_ in S8 Fig.

## Discussion

We are motivated to develop rate models as complementary, or alternative depending on project goals, to detailed spiking network models. We seek frameworks that retain biophysical plausibility, mechanistic interpretability, mathematical tractability and that enable geometrical and dynamical systems viewpoints. Rate models (also referred to as neural mass models) can be computationally efficient and analyzable, providing opportunities to gain insights into local network and multi-area brain dynamics [41,43,44]. In this study we develop and analyze rate models for gamma, of both PING and ING types. The vast literature on modeling gamma oscillations have used spiking networks to highlight key features: the significant role of inhibition, especially its time scales, and reversal potentials, communication delays, architecture, etc. The challenges of robustly coordinating inhibition to produce ING (preventing synaptic inhibition from quenching spiking) have been addressed by introducing delays, gap junctions, etc. [45] (more in the next section).

Our approach to rate-modeling of ING is to extend WC-like models by introducing additional dynamic variables. An *r-s* model without delay does not generate ING. Our novel *r-s-v* and *r-u-s* models capture ING dynamics without using an explicit delay. The models are interpretable mechanistically and we have mathematically analyzed them (e.g., Hopf bifurcations to ING oscillations). We have demonstrated dependence on some parameters comparable to that of a spiking network. Although ING is well-studied in spiking networks, we have used our rate models to provide mean field-like perspectives on major topics in traditional spiking network-based studies of ING such as critical roles of effective delay, hyperpolarizing inhibition versus shunting inhibition, dependence of network frequency on parameters, etc.

Of significance, we have extended our ING-capable firing rate model to an E-I network. To our knowledge, this is the first study of PING and ING in the same network with a rate model. Even in spiking network models, few studies [37] consider the interactions of PING and ING; typically, ING is modeled in isolated I-I network or in E-I network with the excitatory synaptic input is blocked or assumed constant. The firing rate model brings the convenience of visualizing the steady states in the *r*_E_ − *r*_I_ plane that shows how the E and I populations participate in the oscillations as the drive to the I population varies. We revealed a common feature of spiking and rate models in the two-phase recovery between PING episodes.

We also proposed a novel method to generate mean-field time courses and therefore effective bifurcation diagram for spiking network models, which can be applied as a general way for comparisons between spiking network models and firing rate models.

### Synaptic/propagation delay as a critical factor for robust ING

From early spiking network studies [27,46,47], it was learned that ING is fragile to heterogeneity in tonic excitatory drive. Several mechanisms have been incorporated to improve the robustness of ING: (1) Neltner et al. [48] showed that synchrony in heterogeneous inhibitory network was easier at higher firing rates when both the external drive and the coupling strength is strong; (2) Tiesinga & José [49], on the other hand, studied the stochastic weak synchronization of inhibitory network where firing rates are very low and illustrated that synchronization in this regime is robust to cell heterogeneity and synaptic noise; (3) Tikidji-Hamburyan et al. [50] proposed another mechanism of robust ING in a network with Type II cells where the drives to all cells are subthreshold and synchronization is generated by cell resonance and post-inhibitory rebound; (4) Several studies show that explicit synaptic/propagation delay can promote synchrony in both phase-coupled oscillators [51] and an inhibitory spiking network [52]; (5) Paired recording in mice hippocampal slices [53] revealed that the inhibitory synapses are fast and strong with notable conduction delays. Besides chemical synapses, gap junctions also exist. With numerical simulations in an inhibitory network model, the authors confirmed that these factors all promote the network coherence [54,55].

In a firing rate model, some of these synchrony enhancement mechanisms such as cell resonance, distributed delays, gap junctions are difficult to implement. Therefore, in order to generate robust ING, we focused on incorporating delay either implicitly or explicitly between cellular activation and postsynaptic drive.

One approach is to add a pre-synaptic delay, explicit (*r-s* delay model [32]) or implicit (*r-u-s* model), so that the firing rate can develop before being quenched by synaptic activation. Possible biophysical interpretations are propagation delays (fixed as in the *r-s* model or temporally dispersed, represented by *u*(*t*)) or a transmission delay that involves a pre-synaptic first-order process, *u*(*t*). Explicit delay in the *r-s* delay models correspond to case (4) above, in spiking network studies. As reported in [52], the delay duration can significantly influence the network oscillation frequency.

In another approach, we included delay implicitly in the *r-s-v* model (from *s* to *r*) by adding a mean voltage variable as an intermediate temporal integration step between the synaptic field and generation of firing rate. This is a mean-field description akin to the now-classical spiking inhibitory network in the setting of Wang & Buzsáki [27], where no explicit synaptic delay is included but the synaptic gating variable, driven by the pre-synaptic membrane potential, has both rise and decay dynamics We chose the *r-s-v* model to check the dependence of ING on parameters for two reasons: (1) The spiking network is better-studied under this setting; (2) The dynamical equation of mean membrane potential allows the comparison between shunting inhibition and hyperpolarizing inhibition.

Several previous studies (e.g., [27]) found that hyperpolarizing inhibition promotes synchrony, and we find this also for our *r-s-v* model without synaptic delay. However, there is also evidence that shunting inhibition supports more robust synchrony under different network settings. Vida et al. [56] demonstrated that in an inhibitory spiking network with delays and fast synapses, shunting inhibition can not only support gamma oscillation but also improve the oscillation’s robustness. We tested this by adding a *u*-process to our *r-s-v* model so that both voltage- dependent synaptic current and presynaptic delay were represented. We found that, for a fixed input drive, the maximal inhibitory reversal potential increased slightly and then decreased with increasing *τ_u_* (results not shown). We conclude that hyperpolarizing inhibition is still more favored over shunting inhibition even with synaptic delay. Perhaps the inconsistency with the findings of Vida et al. [56] arise since gap junction coupling is not considered in our firing rate models. Tikidji-Hamburyan & Canavier [57] show that shunting inhibition promotes ING in a spiking network of cells with Type I excitability whereas hyperpolarizing inhibition promotes ING in a network of cells with Type II excitability. We are unable to make a comparison just now for a rate model since we have yet to implement Type II cells in our rate models, but this is an interesting future direction.

### Alternate firing and cycle skipping in ING

In three rate models, we found that the ING oscillation emerges and terminates via supercritical Hopf bifurcations as the drive current *I*_I_ passes through critical values. Hence, the firing rate oscillation amplitude increases gradually from zero with *I*_I_ at the onset of oscillations and decreases to zero at the offset. We interpret this low amplitude mean-field oscillation as weakly-modulated spiking, where the rhythm can be seen at the population level, but not all cells participate each cycle.

Sparse firing of I cells was observed in theoretical studies of I-I spiking networks [42,49,58–60]. In an inhibitory spiking network with heterogeneous driving current, we found (but not shown here) that I cells that receive strong drive typically fire more than those with lower drive. That is to say, cells with small input drive skip cycles. We also observed in a network with heterogeneous connectivity that cells with high in-degree skip more cycles than cells with low in-degree tends to fire in each cycle (results not shown).

Clustered patterns of interneuron firing can arise spontaneously among cells firing alternately in simulations at both cellular and network level. In a pair of integrate-and-fire cells with mutual inhibitory coupling, fast-rising synaptic activation generates alternate firing while slow synaptic activation leads to synchrony [61]. In a heterogenous inhibitory network, the cell activity transitions sharply from synchronous to anti-phase firing as the heterogeneity of excitability is increased [46]. In a limited sense, the dynamics of a rate-based model can resemble features of alternate firing and cycle skipping as seen by decreased amplitude and altered occurrence of firing episodes. In simulations with two identical I populations, each modeled by a set of *r-s-v* rate equations we did not find a stable anti-phase pattern; the firing rate oscillation profiles of the two populations were identical and synchronized. However, with stronger intra-connectivity than inter-connectivity, we found examples of bistable activity, alternating cycles of firing episodes.

### PING-ING interactions and relative frequencies

Only a few modeling studies have focused on (or even mention) interactions between PING and ING. Previous studies of ING were usually conducted for an isolated I-I network with tonic input. Here, we found in an E-I spiking network model with an asynchronously active initial state, the network dynamics evolve in one of the three ways: (1) I cells begin to synchronize by the ING mechanism and during a phase of reduced inhibition the E cells break away with an episode of firing that further synchronizes the I cells and PING emerges. (2) I cells synchronize by ING, but the E cells, unable to follow the I cells’ rhythm, remain asynchronous. (3) The nearly constant drive from asynchronous E cells is too weak to recruit the I cells into synchrony so both populations are in asynchrony.

Viriyopase et al. [37] studied the competition and cooperation of PING and ING in an E-I spiking network and found the mechanism that generates higher frequency wins. We observed qualitatively similar behavior in our six-variable E-I firing rate model (Fig 7) and in the spiking E-I network of Hodgkin-Huxley-like units (Fig 1): PING transitions into ING as the drive to I population increases. With strong drive to I cells, the ING frequency in the isolated I-I system is high. The E cells fail to follow high frequency because either (1) The inhibition is too strong, so the E cells are nearly silent, or (2) The network frequency is too fast, so the E cells spike throughout the interval between two I-episodes. In contrast, when the drive to I cells is moderate, the E cells are not overly suppressed and the time between two I-episodes is long enough, allowing for both synaptic inhibition to decay and for E cells to recover. Transitions from PING to faster ING in an E-I spiking network were also reported in [35] for increasing drive to I cells and in [60] by increasing the synaptic conductance between I cells.

In our E-I rate model, the relative difference of PING and ING frequencies depend on the drive to both E and I populations. As illustrated in Fig 7A and 7D, PING is significantly lower in frequency than ING and also compared to PING in the spiking models. While the rate model’s PING frequency is insensitive to the drive in the I population, it depends strongly on the drive to the E population. For example, PING frequency doubles by an increase of 0.2 in *I*_E_ (Fig 7E) and with little change in the ING frequency. For this study we did not fine-tune parameter values (such as time constants) of the E-I firing rate models to achieve strict quantitative agreements with data or spiking models, although this could be done in future work.

We documented the dependence of ING frequency on parameters in the firing rate I-only models. In the *r-s-v* model, network frequency ranged from 30 Hz to 200 Hz, achieved by manipulating only the three time constants (Fig 3). Trivially, by multiplying all time constants by a factor the model’s oscillation frequency is rescaled without changing the phase space rhythmic trajectory. For example, multiplying all time constants by 2 leads to a minimum frequency in the low gamma range (around 45 Hz). Nevertheless, the factor can’t be arbitrarily large or small since *τ_s_* and *τ_v_* are somewhat constrained by biophysical considerations.

In general, our *r-s-v* model shows a relatively high gamma frequency range (around 50-100 Hz when *I*_I_ = 1). The *r-u-s* model and *r-s* delay model with the same external drive, however, cover a lower frequency range (around 30-60 Hz) probably because the delay occurs just one stage presynaptic rather than two stages as in the *r-s-v* model. In the *r-s* delay model, the maximal frequency of oscillation decreases as the duration of the explicit delay increases (S5 Fig), as observed in spiking network models [45,52]. In several spiking network models for ING low frequencies around 30-50 Hz (e.g., [37]), we noticed that explicit synaptic delays were typically included.

### Future directions

We see opportunities for further investigation of rate models of network-based oscillations, some related to relaxing limitations or compromises of these models and others to expanding applications. We directed effort into rate modeling of ING to achieve flexibility in an E-I setting, allowing for the possibility that in some parameter regimes an ING-like rhythm emerges as dominant. Hence in our consideration of limitations, here, we devote several points below to accounting for ING in the rate models.

### Cellular level resonance and Type 2 excitability

It’s known that in spiking networks for ING cellular resonance and rebound provide robustness to the oscillations [50]. Hence, it would be natural to include an intrinsic dynamic recovery variable *w* into, say, the *r-s-v* model that could endow Type II excitability. In this case, both *s* and *w* become negative feedback terms in the balance equation for *v*. We wonder how two negative feedback variables might be helpful for supporting ING and enhancing robustness, especially since our rate model does not have positive feedback or a regenerative process. If spiking (i.e., spike rate) drives synaptic activation, *s*, which variable, *v* or *r*, would effectively drive *w*? What form would represent a mean field description of *w*? We note that gamma oscillations emerge in our rate models by Hopf bifurcations; the network model is of Type 2 although there is no pre-condition for Type 2 at the cell level. The sigmoidal gain function is more reminiscent of Type 1 behavior at the cell level.

### Steep gain function in rate model

We have found that some gain function in the rate models needs to have a steep slope. This feature needs closer investigation. This constraint may be linked to the lack of a regenerative process. A formal derivation of these rate models from, and comparison with, spiking models would likely shed some light here. In rate models, the gain function is typically related to the cellular frequency-current relation which can have a steep rise near the onset, smoothed a bit by the assumed effects of noise and heterogeneity. However, we note that the operating range of ING in our spiking models is not necessarily in this steeply portion of the frequency-current relation. Interestingly, the three-variable rate model for a network of heterogeneous QIF units [34] includes a regenerative term in the dynamical equation for *V*(*t*) and the authors stress the significance of spike upstroke for robustness; there is apparently no need for an additional steepening factor in the model (nor for an explicit delay).

### Significance of delays

While our spiking models did not include propagation delays, many other investigators incorporate delays, especially for modeling ING. We found in the r-s-delay model that having an explicit delay (S5 Fig, S6 Text) can considerably relax the constraint on needing a gain function with a steep slope. This point links to the previous point above, together suggesting that a formal derivation of a rate model could resolve the need for the compromise in implementing WC-like models for ING by incorporation of a steep gain or an explicit delay.

### Derivation of rate model

Some valuable steps have been made with analytic treatments for mean field descriptions in particular parameter ranges. Geisler et al. [62], following on [27], studied the case of a weakly synchronous population rhythm, in the form of sinusoidally-modulated sparse firing probability. The model included transient synaptic conductance and delays (latencies) and described high frequency weak ING oscillations that matched well with spiking network simulations and low frequency weak (classical gamma range) PING oscillations. Such weakly synchronized oscillations we believe are captured by our rate models in which oscillations arise by super-critical Hopf bifurcations (here and in [33]). The weakly modulated I-I oscillation and emergence by Hopf bifurcation was also analyzed using a Fokker-Planck approximation for an LIF network with delay and pulse-coupled inhibition [63]. While these approaches have provided valuable insights, the underlying assumptions and form of solutions carry limitations and both include delays.

We encourage efforts for further analysis and derivations that may reveal how intrinsic properties together with synaptic dynamics, *g*_syn_(*t*), and interactions between units can that lead to rate models that provide meaningful and easily interpretable descriptions. We can hope for rate models that are applicable across the range from weakly modulated sparse firing to more synchronized episodes and across a range of frequencies, that can describe the situation of E-I as well as allowing for potential dominance of I-I. Can we bridge from stochastic dynamics, say, based on the integrate-and-fire family as starting points that lead to low-order MF models and perhaps parameter-tunable based on electrophysiological data? Maybe the WC-framework can be tweaked and extended to overcome some of the current limitations. What form of gain function, formally derivable, is more appropriate than an assumed high-gain sigmoidal-like input-output relation? By developing procedures or guidelines to fit rate-model parameters to simulations of spiking models, one could, hopefully, illustrate how to approximately fit the firing rate model to experimentally recorded data such as cell excitabilities, synaptic input kinetics and connectivity.

### Looking forward

We hope that our results for these idealized extensions of the WC-framework provide options for further applications and extensions of rate models to describe qualitative aspects of E-I local network dynamics. The role of inhibition in balancing and stabilizing excitation is being well-explored with rate models (e.g., [64]). Destabilization of the inhibition-stabilized state and emergence of network oscillations is a natural application (e.g., [65]). In consideration of co-existent slower dynamical mechanisms, such destabilization could describe with rate models, say, gamma oscillations nested within a theta rhythm (e.g., [66,67]). Given the existence of numerous inhibitory subtypes, the rate modeling framework can be used to explore possibilities for oscillations, including disinhibition, and computation more generally. The rate model framework can incorporate dynamical processes, intrinsic cellular or synaptic, that operate on the timescale of a few to several spikes and, say, modulate gamma or other rhythms. Synaptic depression and facilitation, spike frequency adaptation, slow ionic currents have been incorporated into rate models (e.g., [68]). Parameter variation and bifurcation analyses with rate models can be effective in probing network dynamics. They provide exploratory opportunities for addressing questions and situations in which some biophysical details of spiking may be put aside.

## Methods

## 1 Firing rate models, parameters, and numerical simulations

### 1.1 Three-variable r-s-v model for the I-I network

The three-variable rate model describes the dynamics of firing rate *r*, synaptic gating variable *s*, normalized membrane potential *v*. The variables *r* and *s* are both between 0 and 1; *v* is normalized such that the *v* = 0 represents rest and *v* = 1 represents 0 mV for the excitatory synaptic reversal potential. The equations are

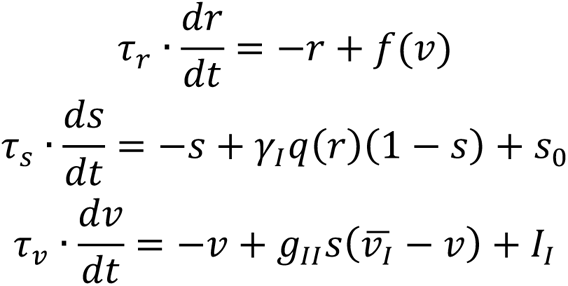

where the dependence of steady state firing rate on the membrane potential is

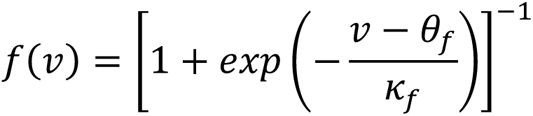

and dependence of synaptic activation rate on the firing rate is

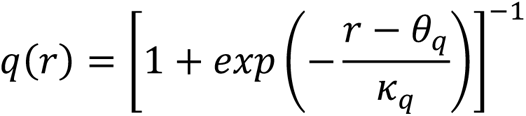

They are both sigmoid increasing functions with values between 0 and 1. Each is centered at *θ* and the slope at the point (*θ*, 1/2) is 1/(4*κ*); small *κ* means steep input-output function.

The model’s control parameter values corresponding to Fig 2B top are listed in Table 1. For the bifurcation diagrams, all parameters keep their values as in the table except for the bifurcation parameter that is varied in a certain range.

**Table 1.** Parameters for three-variable *r-s-v* model for the I-I network.

|  |  |  |  |  |
| --- | --- | --- | --- | --- |
| Parameter | $g_{II}$ | $\gamma_I$ | $I_I$ | $s_0$ |
| Value | 5 | 3 | 1 | 0.05 |
| Parameter | $\theta_f$ | $\kappa_f$ | $\theta_q$ | $\kappa_q$ |
| Value | 0.4 | 0.01 | 0.5 | 0.1 |
| Parameter | $\tau_r$ | $\tau_s$ | $\tau_v$ | $\bar{v}_I$ |
| Value | 2 ms | 10 ms | 5 ms | -0.1 |

### 1.2 Two-variable r-s delay model for the I-I network

The two-variable *r-s* delay model was adapted from [32]. Parameter values were modified to be the same as the values in the *r-s-v* model and *r-u-s* model for comparison.

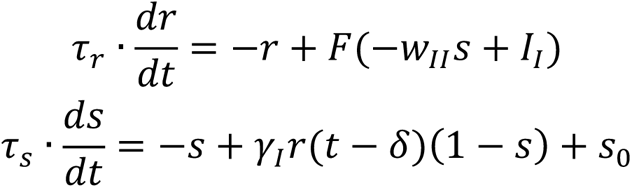

The input-output function takes the same form with the *r-u-s* model except for different parameter values of *θ_F_* and *κ_F_*.

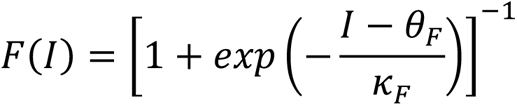

The control parameters that generate the time course in Fig 2C are listed in Table 3.

### 1.3 Three-variable r-u-s model for the I-I network

We introduced an alternative way to include an implicit delay in Fig 5. The model’s equations are

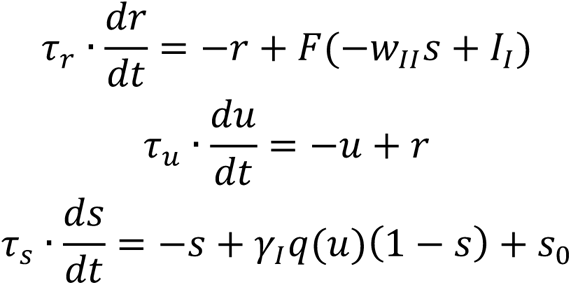

The input-output function is

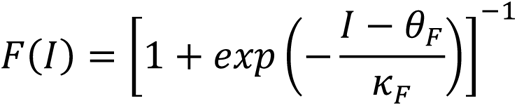

As for the *r-s-v* model, the scaling factor of synaptic activation rate is

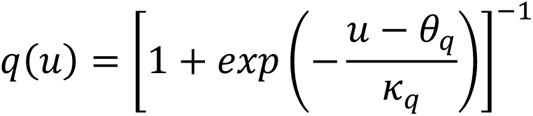

The control parameters that generate the time course in Fig 5B are listed in Table 2. The values were chosen to be the same as those for the *r-s-v* model when the corresponding parameters are shared in the two models. All parameters were fixed except for the inverse steepness *κ*_F_ and the external drive *I_I_* in the bifurcation diagrams.

**Table 2.** Parameters for two-variable r-s delay model for the I-I network.

| Parameter | $w_{II}$ | $\gamma_I$ | $I_I$ | $\theta_F$ | $\kappa_F$ | $s_0$ | $\tau_r$ | $\tau_s$ | $\delta$ |
| --- | --- | --- | --- | --- | --- | --- | --- | --- | --- |
| Value | 5 | 3 | 2 | 0.5 | 0.01 | 0.05 | 2 ms | 10 ms | 1 ms |

**Table 3.** Parameters for three-variable *r-u-s* model for the I-I network.

| Parameter | $w_{II}$ | $\gamma_I$ | $I_I$ | $s_0$ |
| --- | --- | --- | --- | --- |
| Value | 5 | 3 | 2 | 0.05 |
| Parameter | $\theta_F$ | $\kappa_F$ | $\theta_q$ | $\kappa_q$ |
| Value | 0.4 | 0.05 | 0.5 | 0.1 |
| Parameter | $\tau_r$ | $\tau_u$ | $\tau_s$ | |
| Value | 2 ms | 5 ms | 10 ms |  |

### 1.4 Six-variable r-s-v model for E-I network

To model the interaction between E and I populations, we included three equations for the E population with three-variable *r-s-v* rate model for the I population, as well as additional synaptic current terms to the voltage equation. The equations for the six-variable model are

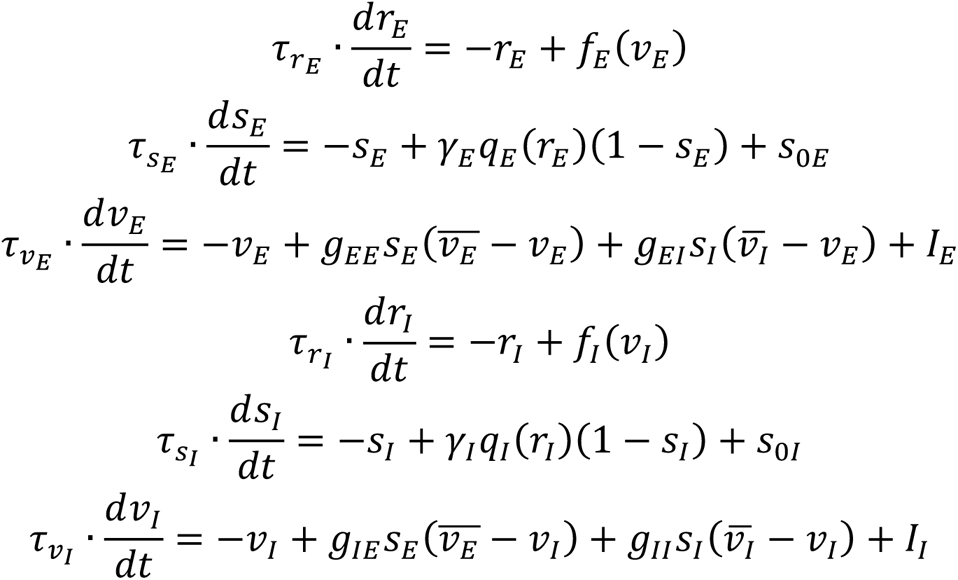

The E and I populations have different parameters for the input-output function, where the function for I population is steeper than E population.

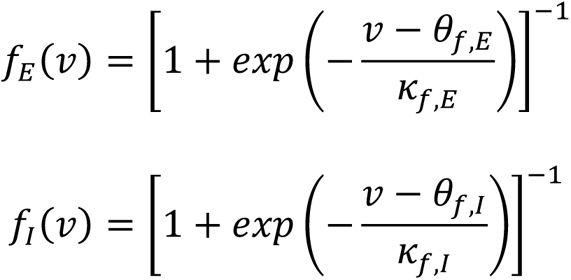

The synaptic activation rate is the same for the E and I populations in simulations shown in this paper.

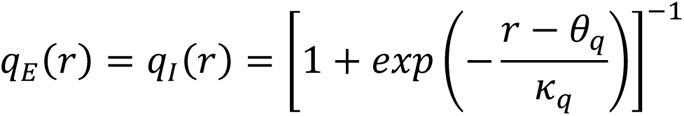

The PING time course in Fig 7A corresponds to the control parameter values of Table 4. Replacing *I*_I_ by 0.5 will give the ING time course in Fig 7B. *I*_E_ = − 0.2 in Fig 7A-D, *I*_E_ = 0 in Fig 7E.

**Table 4.** Parameters for six-variable *r-s-v* model for the E-I network.

|  |  |  |  |  |  |  |  |  |
| --- | --- | --- | --- | --- | --- | --- | --- | --- |
| Parameter | $g_{EE}$ | $g_{EI}$ | $g_{IE}$ | $g_{II}$ | $\overline{v_E}$ | $\overline{v_I}$ | $I_E$ | $I_I$ |
| Value | 2 | 6 | 2 | 2 | 1 | -0.1 | -0.2 | 0 |
| Parameter | $\theta_{f,E}$ | $\kappa_{f,E}$ | $\theta_{f,I}$ | $\kappa_{f,I}$ | $\theta_q$ | $\kappa_q$ | $\gamma_E$ | $\gamma_I$ |
| Value | 0.2 | 0.05 | 0.4 | 0.1 | 0.5 | 0.1 | 3 | 3 |
| Parameter | $\tau_{rE}$ | $\tau_{sE}$ | $\tau_{vE}$ | $\tau_{rI}$ | $\tau_{sI}$ | $\tau_{vI}$ | $s_{0E}$ | $s_{0I}$ |
| Value | 1 ms | 3 ms | 3 ms | 2 ms | 10 ms | 5 ms | 0.2 | 0.05 |

### 1.5 *r*E and *r*I - steady state relationships (SSRs) for the six-variable *r-s-v* model

To obtain the *r*_E_ and *r*_I_ - SSRs, we first set each equation in the system into steady state. Then we write *v*_E_ as a function of *r*_E_ using d*r*_E_/d*t* = 0 and *s*_E_ as a function of *r*_E_ using d*s*_E_/d*t* = 0. The relations are given by

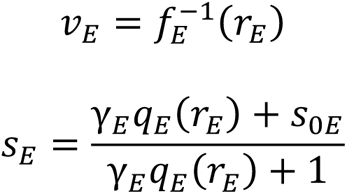

Similarly, we write *s*_I_, *v*_I_ as functions of *r*_I_.

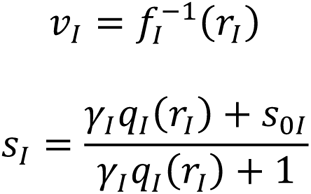

In the equation d*v*_E_/d*t* = 0, by replacing *v*_E_, *s*_E_ and *s*_I_ with the above functions of *r*_E_ and *r*_I_, we obtain an implicit function of *r*_E_ and *r*_I_ given by

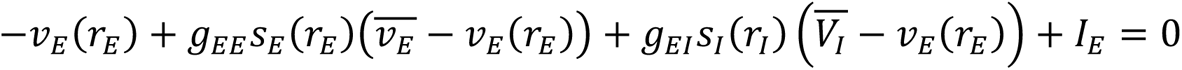

We plot it in the *r*_E_ − *r*_I_ plane as the *r*_E_ - SSR numerically. Similar process is applied to the equation d*v*_I_/d*t* = 0 for the *r*_I_ - SSR.

### 1.6 Numerical Methods

The time courses and one-parameter bifurcation diagrams for the firing rate models were computed using XPPAUT [69]. The data were exported from XPPAUT and imported to MATLAB using Ting-Hao Hsu’s MATLAB Interface for XPP [70] to generate the plots. The numerical integration scheme used in XPP is 4^th^ order Runge-Kutta with time step *dt* = 0.05 ms. Halving *dt* produced no noticeable difference in time courses.

To generate the two-parameter bifurcation diagrams (Fig 3, 4, 6), we first solved for the Hopf boundary (black solid curves). For each fixed parameter #1 (values along the horizontal axis, time constants in Fig 3, *γ*_I_, *g*_II_ and *v*2_#_ in Fig 4, *κ* in Fig 6), we solved for the Hopf bifurcations (denote *I*_HB1_ and *I*_HB2_ for oscillation onset and offset respectively) with respect to parameter #2 (*I*_I_ along the vertical axis) with the MATLAB nonlinear root-finder. The nonlinear system consists of five analytically derived equations. Three of the equations are from the fixed-point condition (the derivatives of three variables, with respect to time are zero); the other two are from the real and imaginary parts of the Hopf bifurcation condition (the eigenvalues of the Jacobian matrix are purely imaginary). The corresponding five unknowns are the three variables, *I*_I_ and the imaginary part of the eigenvalue. With five nonlinear equations and five unknowns, we solved for the values of *I*_I_ at the Hopf bifurcations with judicious choice of initial guess. The coordinates of Hopf bifurcation points were connected by line segments to get a Hopf boundary for oscillation. For each pair of (discretized) parameter values inside the boundary, we ran time courses, calculated oscillation frequency and visualized the frequency in color.

## 2 Linear stability analysis of the standard Wilson-Cowan model

We consider the following version of the Wilson-Cowan model for the I-I network and show that no oscillations can be generated for any choice of parameters.

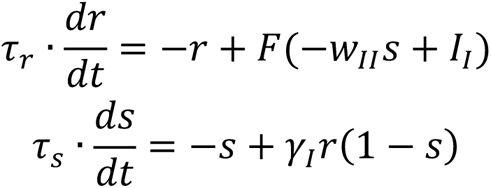

To study the stability of the fixed point, we only need to check the eigenvalues of the Jacobian matrix at the fixed point. The Jacobian matrix is

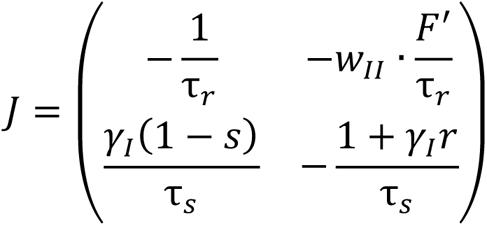

Therefore, the eigenvalues satisfy

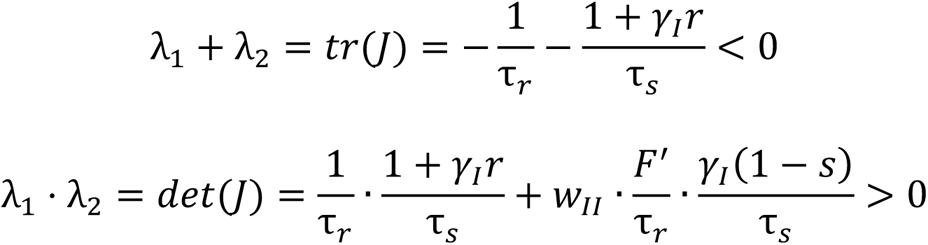

That is to say, the eigenvalues have negative real parts. So, the fixed point is stable and the system can’t generate stable oscillation.

## 3 Linear stability analyses of the three-variable *r-s-v* model

### 3.1 Fixed point and first order approximation near the fixed point

Denote the fixed point of the three-variable system as (*r*\*, *s*\*, *v*\*). Setting the right-hand side of the equations to zero gives a system of nonlinear equations for a fixed point.

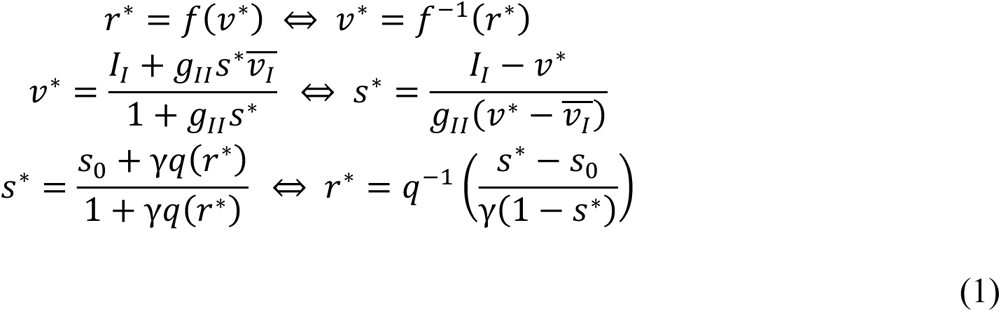

To study the local dynamics near the fixed point, we only need to consider the Jacobian matrix *J* at (*r*\*, *s*\*, *v*\*), which is

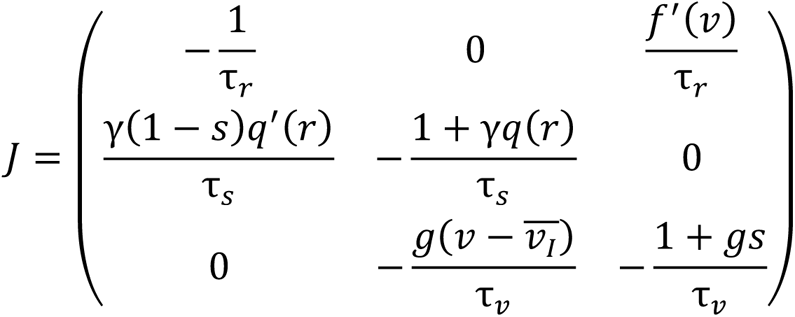

For simplicity of notation, some subscripts (*I* for inhibition) and superscripts (asterisk for fixed point) are omitted from now on. The eigenvalues *λ* of the Jacobian matrix *J* satisfy det(*J* − *λ*·*Id*) = 0 where *Id* is the identity matrix, i.e.

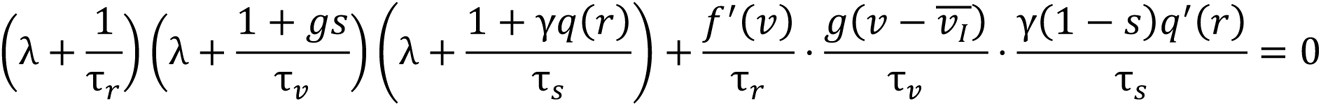

Graphically, the *λ*-values can be interpreted as the roots of a vertically shifted cubic curve, since the first term is a cubic term with 3 negative real roots and the second term is positive. By plotting the left-hand side as a function of *λ* and checking the number of intersections with the zero-line, it can be seen that the shifted-up cubic curve has 3 negative roots if the shift-up term is small; it has 1 negative root and a pair of complex roots if the shift-up term is large. Therefore, the only way for the fixed point of the system to exchange stability is via Hopf bifurcation.

### 3.2 Condition for Hopf bifurcation

At a Hopf bifurcation, the pair of complex eigenvalues *α ± βi* is purely imaginary. Thus, to obtain the condition for the existence of Hopf bifurcation, we substitute *λ* = *βi* and get

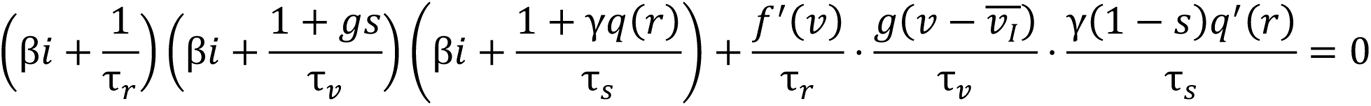

Since the real part and the imaginary part of the left-hand side are both zero, we have

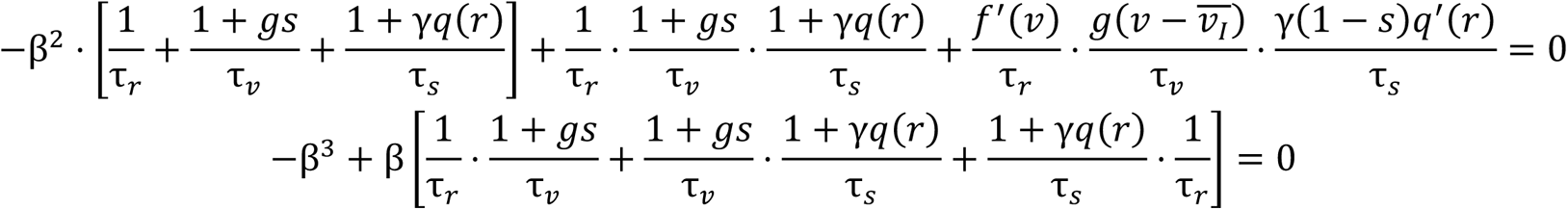

It can be seen from the first equation that *β* ≠ *0*, then the equations for the real part and the imaginary part give two formulas for *β^2^*,

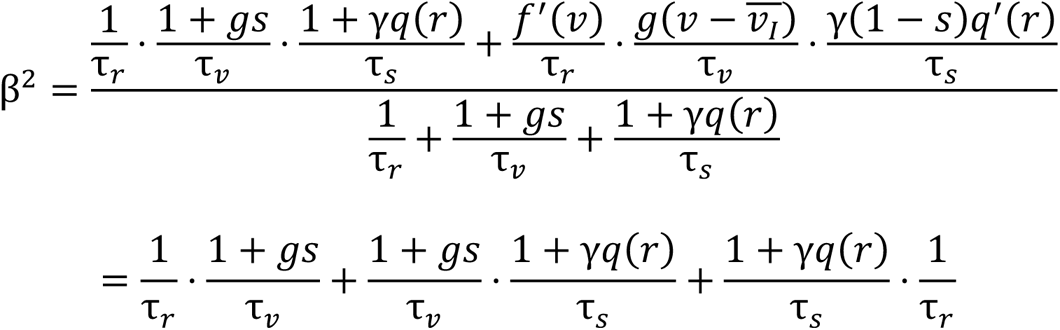

Rewriting the equation yields

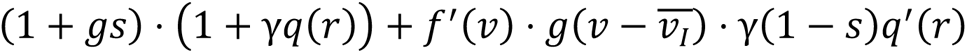

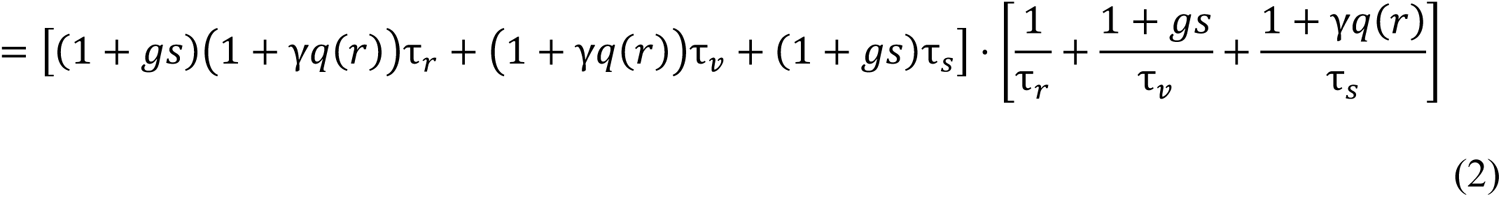

This is a condition for a Hopf bifurcation. With all other parameters fixed, the value for the parameter of interest at a Hopf bifurcation can be solved from the system of nonlinear equations given by the fixed point condition Eq 1 and Hopf bifurcation condition Eq 2. There are 4 equations and 4 unknowns: three variables *r*, *s*, *v* and one parameter of interest such as *I*.

Denote 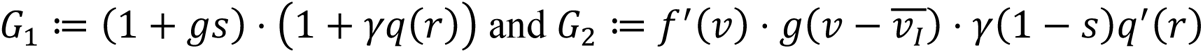 for the two terms on the left-hand side of Eq 2. Applying the Cauchy-Schwarz inequality (see e.g., [71]) to the right-hand side yields

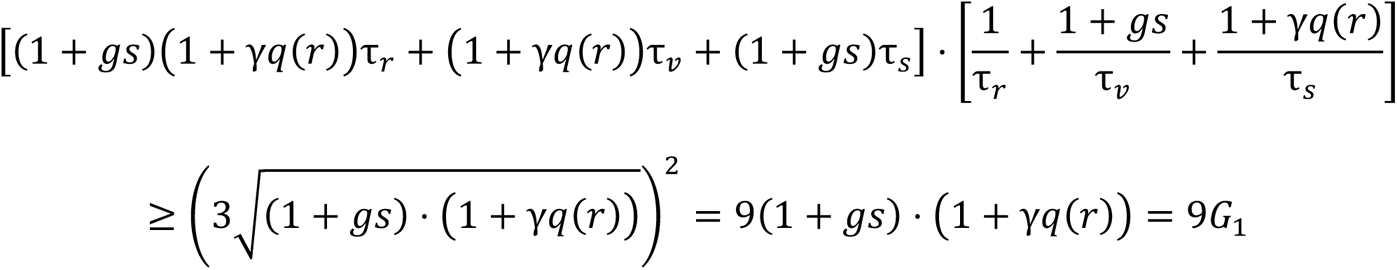

Therefore, a necessary condition for the existence of the Hopf bifurcation is *G*_2_ ≥ *8G*_1_, i.e.

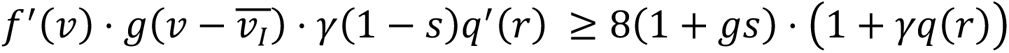

The equality of Cauchy-Schwarz is attained when

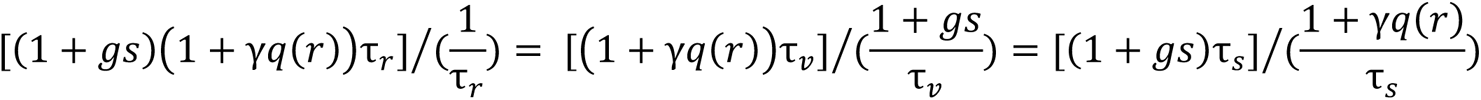

This can be rewritten as

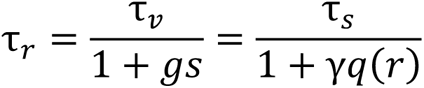

which means the effective time constant of the three variables are the same.

### 3.3 Dependence of the existence of Hopf bifurcation on parameters

In order to have a stable periodic solution, the existence of a Hopf bifurcation is necessary. Therefore, the minimum of the right-hand side of Eq 2 given by Cauchy-Schwarz inequality needs to be smaller than the left-hand side. That is, the three effective time constants being close to each other is a condition that favors the existence of oscillation. Suppose one of the time constants *τ_r_*, *τ_s_*, *τ_v_* is very small (goes to 0) or very large (goes to ∞), then the right-hand side of Eq 2 will go to ∞ and there can be no solution for Hopf bifurcation.

Dependence on the other parameters needs more careful consideration. Recall that the Hopf bifurcation is a fixed point, so it satisfies the fixed point condition Eq 1. By writing *r* and *s* as a function of *v*, *G*_1_ and *G*_2_ in the necessary condition can be rewritten in terms of only one variable *v* and the parameters,

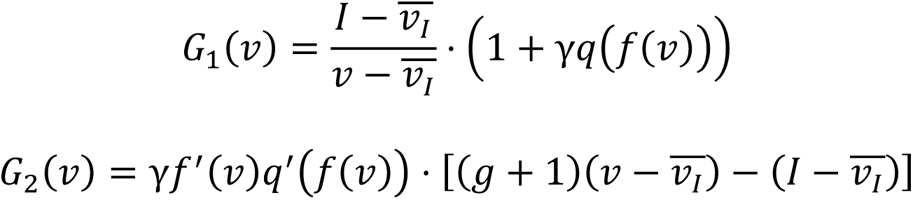

Then *G*_2_ ≥ *8G*_1_ becomes

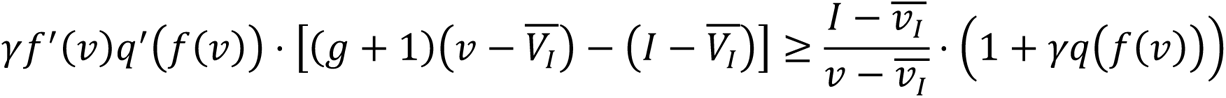

Notice that for a sigmoid function

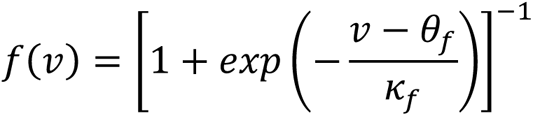

Its derivative is given by

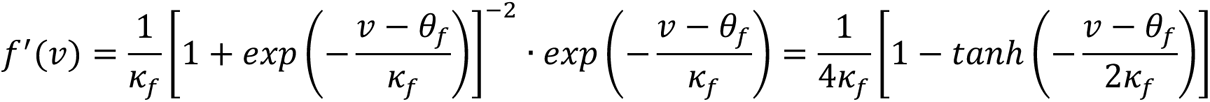

So the derivative at *v* = *θ_f_* (such that *f*(*v*) = 1/2) is *f* ′(*v*) = 1/(4*κ_f_*).

The inequality suggests that to have an oscillatory solution, the input-output function *f*(*v*) needs to be steep (*κ_f_* needs to be small). The conductance *g* has a lower bound and the external drive *I* has an upper bound to ensure the term 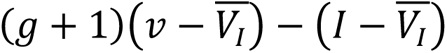 to be positive. Moreover, the synaptic activation rate *γ* has a lower bound since *γ* = 0 sets the left-hand side to 0 and make the inequality fail. These observations match with the numerical bifurcation plots. However, since they are derived from a necessary condition, these constraints are not sufficient to guarantee the existence of Hopf bifurcation. For example, the upper bound for *γ* and *g*, lower bound for *I* in the numerical results are not predicted in this analysis.

## 4 Spiking network models

### 4.1 Hodgkin-Huxley like model for E-I network

The E-I spiking network in Fig 1 consists of 800 E cells and 200 I cells. The single cell model was adapted from [40]. The model equations are

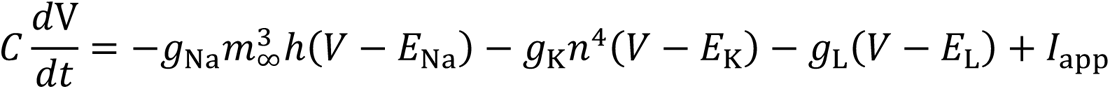

where *C* = 1μF/cm^2^. The maximal conductance of the ionic currents is *g*_Na_ = 24 mS/cm^2^, *g*_K_ = 3 mS/cm^2^, *g*_L_ = 0.02 mS/cm^2^. The reversal potentials are *E*_Na_ = 55 mV, *E*_K_ = −90 mV, *E*_L_ = −60 mV. The applied currents *I*_app_ are heterogenous in both E and I cells. For each E cell, it is drawn randomly from a uniform distribution on [0.38, 0.42] μA/cm^2^ so that the range of the intrinsic frequency of the E cells is [38.03, 40.45] Hz. For each I cell, it is drawn randomly from a uniform distribution on [−1.05, −0.95] μA/cm^2^ (below threshold) for PING and [0.95, 1.05] μA/cm^2^ (intrinsic frequency in [63.32, 67.19] Hz) for ING. The unitless gating variables *m*_∞_, *h*, *n* are given by

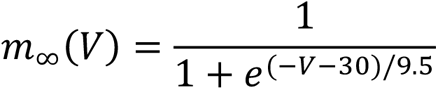

and

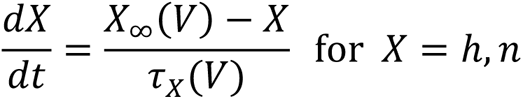

Where

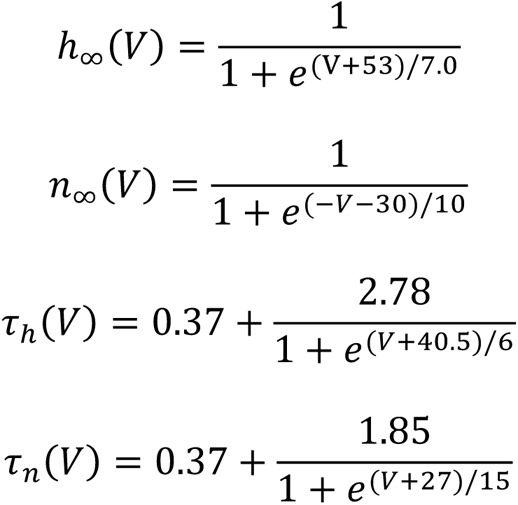

The synaptic current is modeled by

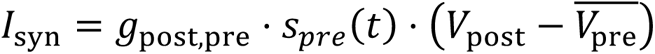

where the maximal synaptic conductance *g*_EE_ = 0.004 mS/cm^2^, *g*_II_ = 0.016 mS/cm^2^, *g*_IE_ = 0.002 mS/cm^2^, *g*_EI_ = 0.004 mS/cm^2^ and the reversal potential 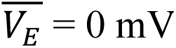, 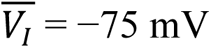. The synaptic gating variable *s* is driven by presynaptic membrane potential. It follows the dynamical equation

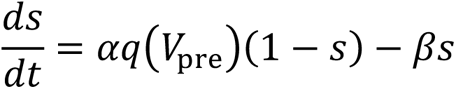

where *q*(*V*) = 1 for 0.5 ms after *V* crosses 0 mV from below and *q*(*V*) = 0 otherwise. The rising rates are *α*_E_ = 14/3 ms^−1^, *α*_I_ = 53/11 ms^−1^, and the decay rates are *β*_E_ = 1/2 ms^−1^, *β*_I_ = 2/11 ms^−1^. The connectivity between neurons was randomly assigned, with connection probability *p*_post,pre_ is *p*_EE_ = 0.05, *p*_EI_ = *p*_IE_ = *p*_II_ = 0.3.

### 4.2 Wang-Buzsáki model for I-I network

The I-I spiking network in Fig 2 consists of 200 I cells. The single cell model and parameters follow Eq 2.1 in [27] and the synaptic dynamics follows Eq 2.4 in the same paper. Each cell has 60 randomly chosen presynaptic cells. The applied current to the cells is heterogenous, following a Gaussian distribution with a mean value *I*_I_ = 2 μA/cm^2^ and standard deviation 0.02 μA/cm^2^. The intrinsic frequencies for 1.98 μA/cm^2^ and 2.02 μA/cm^2^ are 101.04 Hz and 102.52 Hz respectively. Each simulation was run for *T* = 500 ms, in MATLAB using 4^th^ order Runge-Kutta with time step *dt* = 0.05 ms.

### 4.3 Mean field quantities from spiking network simulations for comparison with rate model time courses

To generate analog time courses for the spiking model to compare with time courses of *r*, *v* and *s* from the rate model, we used the averaged synaptic gating variable over cells <*S*> as the population synaptic gating variable *s*(*t*). For the voltage variable *v*(*t*), we computed the average membrane potential <*V*> and scaled it such that 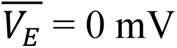 and 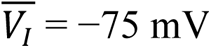 becomes 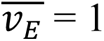 and 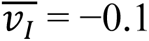 as in the rate model. That is, *v* = 1.1<*V*>/75 + 1. To calculate the instantaneous firing rate *r*(*t*), we counted the number of spikes *N*_spk_ in a sliding window of size 1 ms or 5 ms (indicated in Figure legend) and step 0.05 ms, and calculated R= *N*_spk_/(*N*_cells_·1 ms) = *N*_spk_/(200·10^−3^ s). To ensure the firing rate is between 0 and 1, we normalized the instantaneous firing rate by setting *r* = *R*/*R*_max_ where *R*_max_ is the maximum of the instantaneous firing rate *r*(*t*) over time.

To obtain an analog of the one-parameter bifurcation plot, we generate, as in the previous paragraph, a “mean field” time course from a spiking model run for each parameter value. The temporal average over the run of spiking quantities were also computed for plotting in the bifurcation diagram in lieu of a steady state fixed point (the unstable asynchronous firing state) for the spiking network. We also computed the synchrony measure ranged between 0 and 1 [46] and set 0.5 as a threshold. We distinguished the ING-like pattern (synchrony measure > 0.5) and asynchronous firing (synchrony measure < 0.5) by marking the temporal mean with red and orange points respectively. For the ING case, the maximum and minimum values of the variable on the stable periodic orbit were approximated by those in the analog time course; the oscillation period was approximated by the mean distance between peaks in the ‘time course’ and the oscillation frequency is the reciprocal of period. The ‘bifurcation plot’ in Fig 2A middle and bottom were averaged over 5 trials for each *I*_I_.

## Acknowledgements

J Rinzel acknowledges helpful discussions with LieJung Shiau on rate modeling of ING.

## Supporting Information

**S1 Fig.**
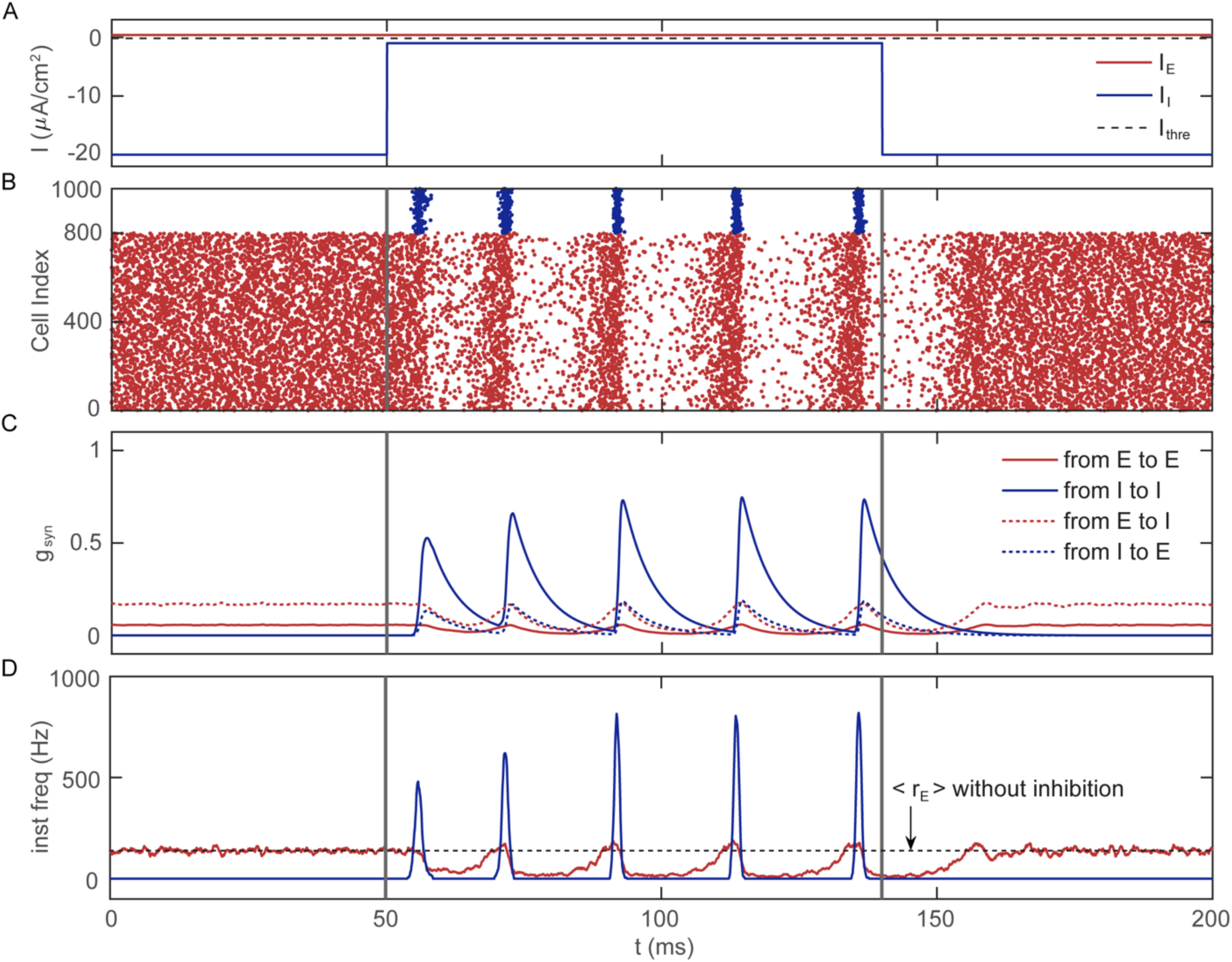
Transient phase of PING. **(A)** E cells receive constant suprathreshold external drive (*I*_E_ = 0.4 μA/cm^2^, red lines), while the I cells receive a subthreshold step drive (*I*_I_ = −1 μA/cm^2^ from 50 ms to 140 ms on a background of suppressive drive *I*_I_ = −20 μA/cm^2^, blue lines). The threshold *I*_app_ = −0.18 μA/cm^2^ is indicated by the black dashed line. **(B)** Raster plot of the E (red) and I (blue) cells shows that the E cells are asynchronous, and I cells are silent in the first 50 ms; when the suppression to I cells is over-ridden at 50 ms (left vertical grey bar), the I cells wait for around 5-10 ms to fire together. The synchronous firing of I cells shuts down the E cells and generates the first PING cycle. As the inhibitory synaptic current decays, the E cells start to fire together and I cells become more synchronized. Temporal summation of successive inhibitory synaptic fields lead to an adaptation to longer PING period. At 140 ms (right vertical grey bar), the I cells are suppressed again, it takes almost the same time as in PING for the E-cell activity to gradually build-up and return to asynchrony. **(C)** Synaptic conductance versus time shows that the recovery of E-cell activity consists of an inhibitory synaptic field decay phase and an excitatory activity build-up phase, each lasting for around 10 ms. **(D)** Instantaneous frequency (window size = 1 ms) of E (red) and I (blue) cells show that the I cells become more synchronized during the adaptation of PING. The mean frequency of asynchronous E-cell activity in absence of I cells shown by the black dashed line indicates that this frequency is the target for E-cell activity in PING.

**S2 Fig.**
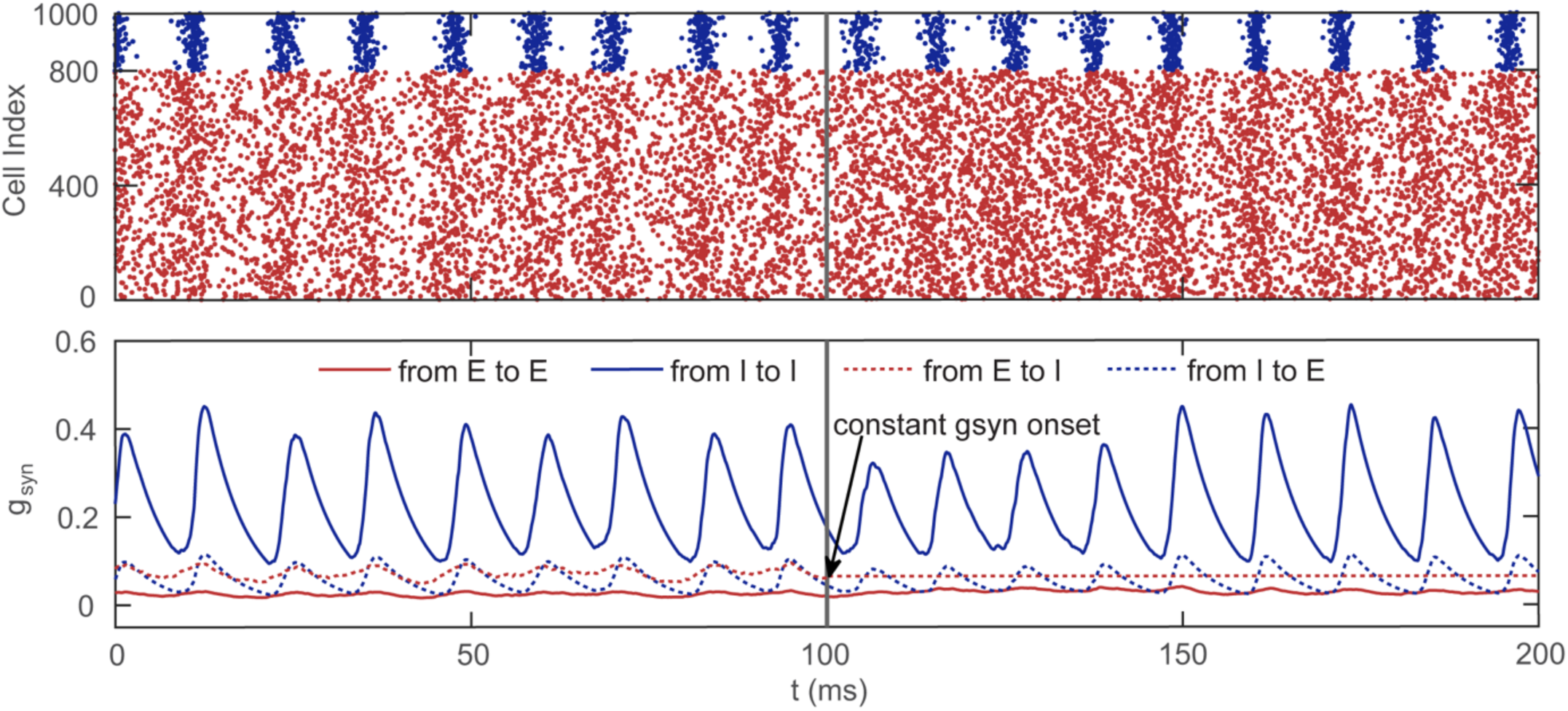
Replacing *g*_syn,IE_(*t*) by a constant (= 0.65 nS/cm^2^) does not noticeably change the ING oscillation. The raster plot in the top panel shows that in the first 100 ms, the E-I network generates ING oscillation (*I*_E_ = 0.4 μA/cm^2^ and *I*_I_ = 1 μA/cm^2^); in the second 100 ms, the synaptic current from E to I (red dashed curve in the bottom panel) is replaced by a constant value and the ING oscillation persists, indicating the tiny oscillation in *g*_syn,IE_(*t*) does not contribute to the synchronization of I cells.

**S3 Fig.**
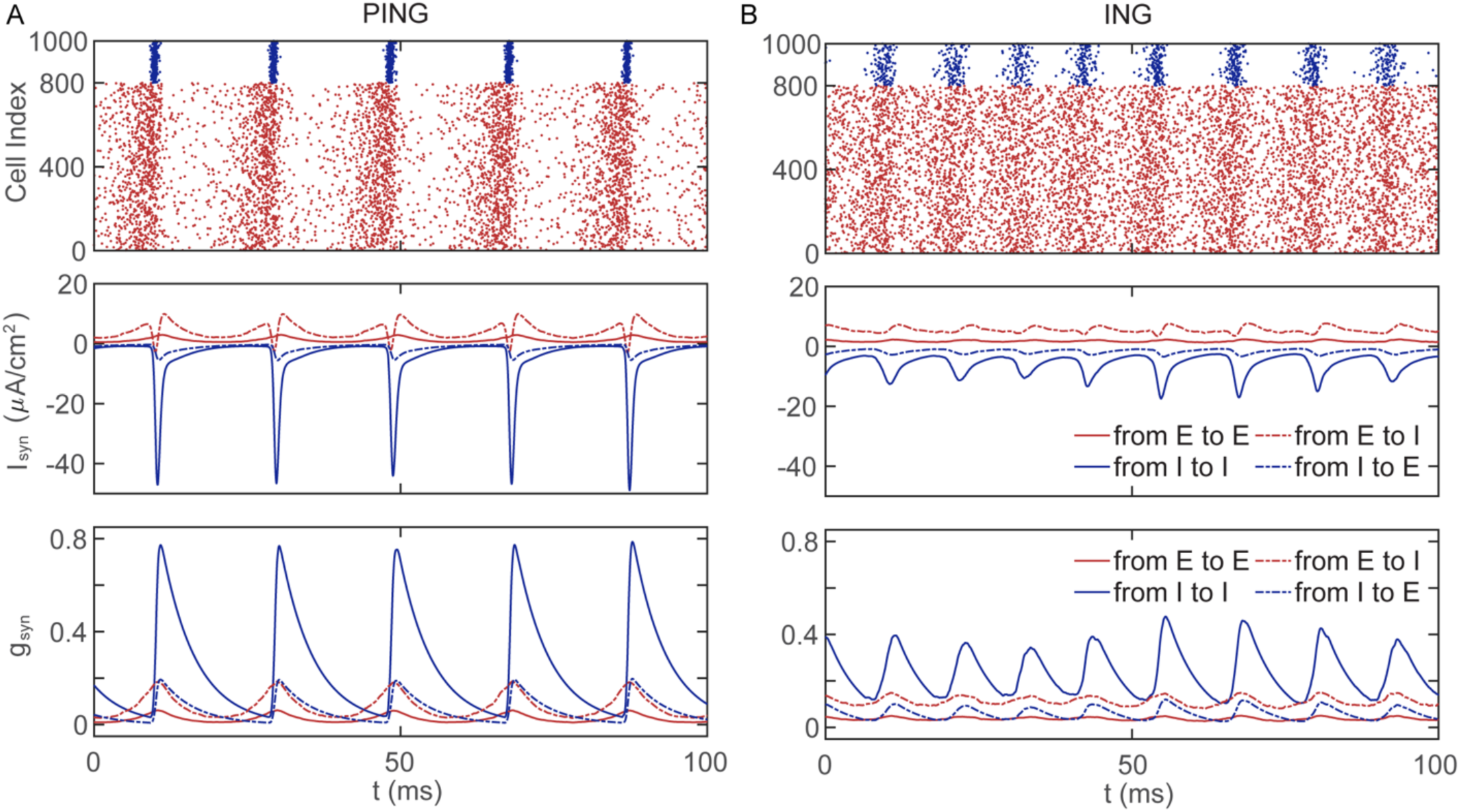
An example realization of PING/ING bistability in the spiking network. **(A)** When *I*_E_ = 0.4 μA/cm^2^ and *I*_I_ = −0.8 μA/cm^2^, the network generates a PING oscillation of 50 Hz when initialized synchronously. **(B)** With the same external drives to E and I cells (same realization of heterogeneity in external drive) and the same realization of network structure, the network generates an ING oscillation of 80 Hz when initialized asynchronously.

**S4 Fig.**
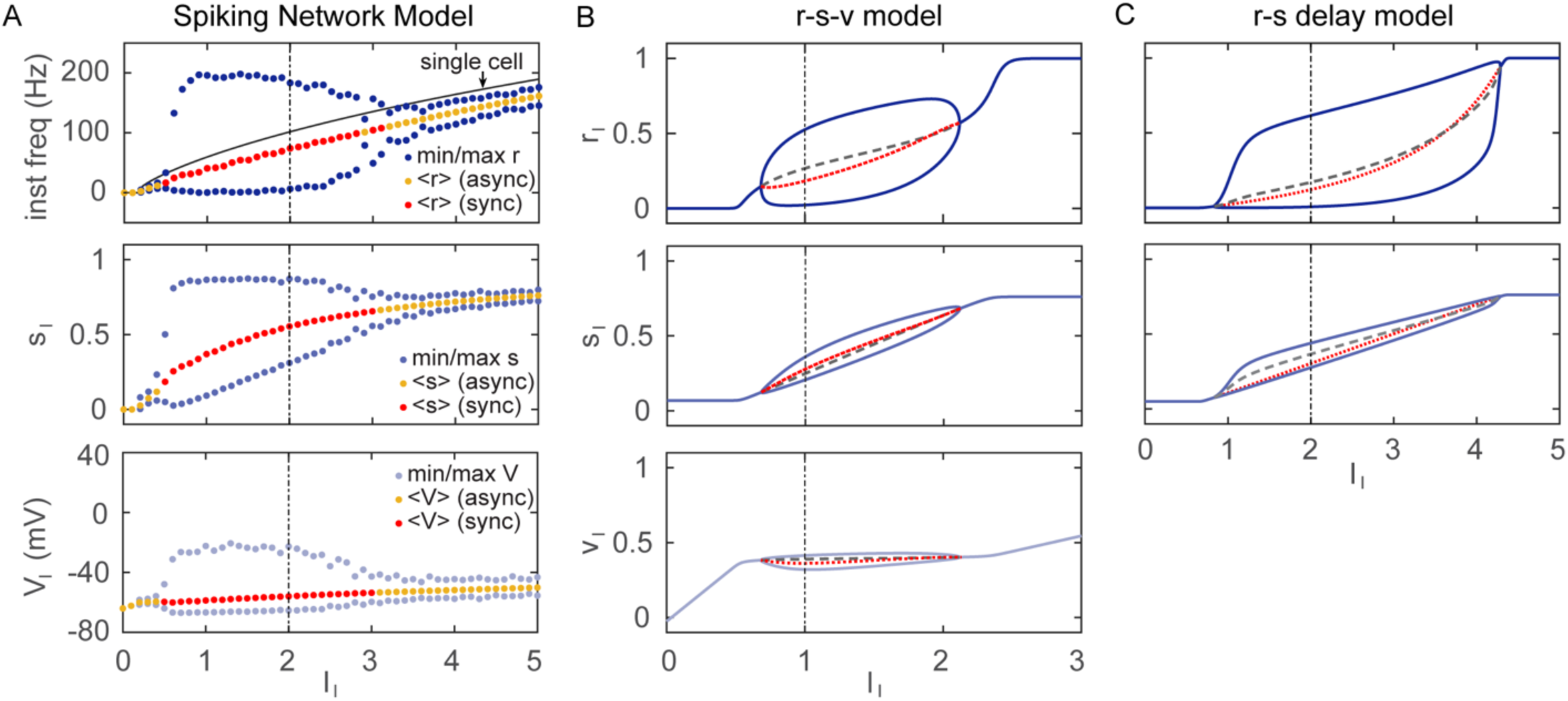
Bifurcation diagrams of the *r-s-v* model (B) and r-s delay model (C) and analogous diagram for the spiking network model (A) with input drive *I*_I_ as the control parameter. Line color and type as in Figure 2 (main text). The middle panels show min/max and mean values for synaptic gating variable *s* and bottom panels (for A and B) show mean membrane potential *v*; the stable/unstable fixed points are shown for the rate models. Notice that the mean membrane potential is not normalized for the spiking network model and the oscillation amplitude reflects the level of synchrony. The flat dependence of the unstable fixed point (in B, lower panel) of *v* on *I*_I_ near *θ_f_* = 0.4 is due to the steepness of the input-output function *f*(*v*).

**S5 Fig.**
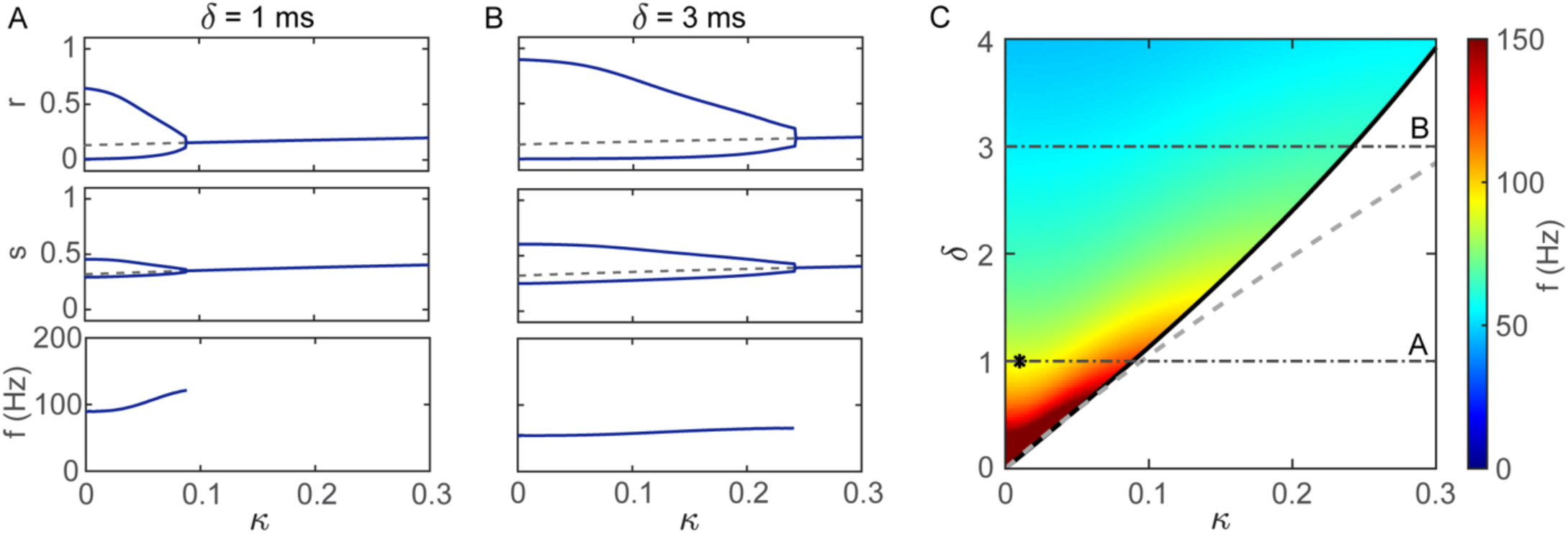
Oscillation in the *r-s* delay model does not rely on steep input-output function. **(A)** Bifurcation diagram for the inverse steepness *κ* when the explicit delay *δ* = 1ms. The Hopf bifurcation (*κ*_HB_ = 0.088) representing the maximum *κ* for oscillation is greater than that in *r-s-v* model (*κ*_HB_ = 0.026). **(B)** Same with **A** but for *δ* = 3 ms. The value of Hopf bifurcation (*κ*_HB_ = 0.241) is greatly increased and the oscillation frequency is decreased from *δ* = 1 ms. **(C)** Two-parameter bifurcation diagram for the inverse steepness *κ* and the explicit delay *δ* with *I*_I_ fixed (*I*_I_ = 2). Horizontal dotted lines corresponds to **A** and **B**; the asterisk shows the parameter values for the time course in Fig 2C (*δ* = 1 ms, *κ* = 0.01). As the explicit delay increases, less steepness is required for oscillation. The implicit delay sets an upper bound of oscillation frequency. The frequency decreases with increased *δ* and has weaker dependence on *κ*. The linear approximation in S6 Text (grey dashed curve) well approximates the Hopf bifurcations of the *r-s* delay model (black solid curve) when *δ* is small.

### S6 Text. Analysis on the effect of the explicit delay in the *r-s* delay model.

Consider the *r-s* delay model and expand the delay term to the first order, i.e. *r*(*t*−*δ*) = *r*(*t*) – *δr*′(*t*) *+* O(*δ^2^*). Then an approximation of d*s*/d*t* equation is

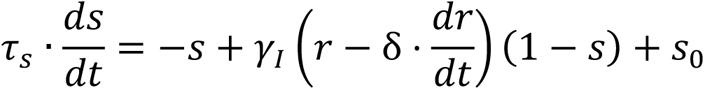

Substituting the d*r*/d*t* term by the corresponding ODE yields

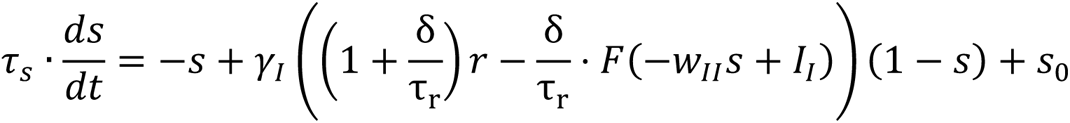

Therefore, an approximation of the r-s delay model can be written as

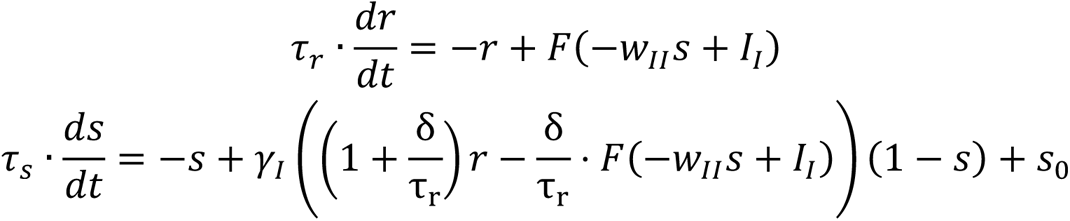

Notice that at the steady state, *r = F*(−*w*_II_*s + I*_I_). The Jacobian matrix at the fixed point can be simplified to

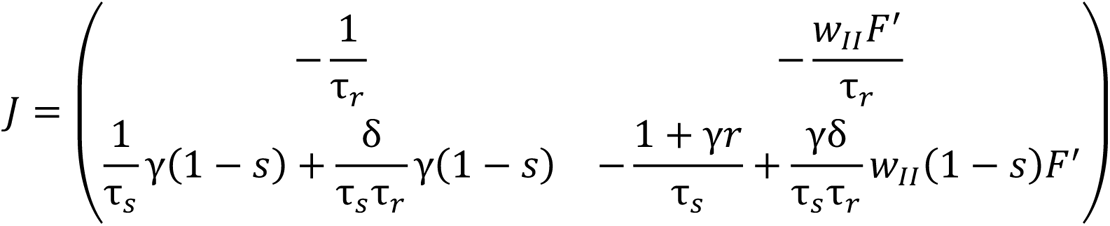

It follows that

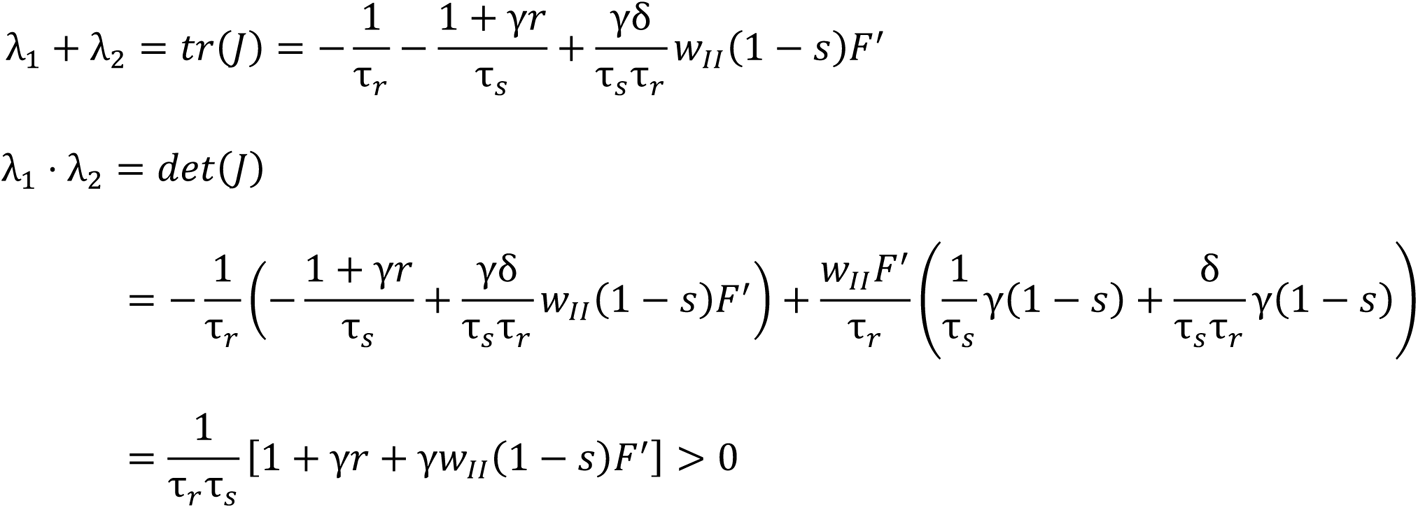

Therefore, the oscillation can only emerge from Hopf bifurcation as *λ*_1_*+ λ*_2_ crosses 0. Notice that at the fixed point,

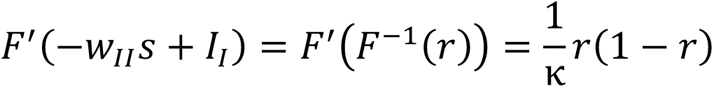

So the only positive term in the *λ*_1_*+ λ*_2_ equation can be rewritten as

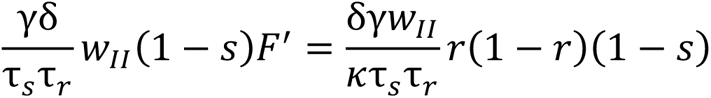

By setting *λ =± βi*, the imaginary part (*λ*_1_*+ λ*_2_ = 0) gives the condition for Hopf bifurcations (grey dashed curve in S5 Fig).

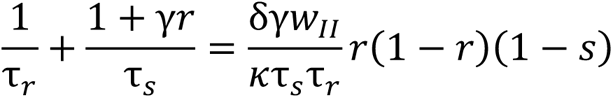

It can be seen that in order for the Hopf bifurcation to exist, *δ* should be large and *κ* should be small. Since *κ* has small influence on the steady state values of *r* and *s*, *δ* and *κ* scale almost linearly on the curve of Hopf bifurcations. Increasing *δ* will allow more flexibility in *κ*. The real part gives a formula for oscillation frequency at the Hopf bifurcations.

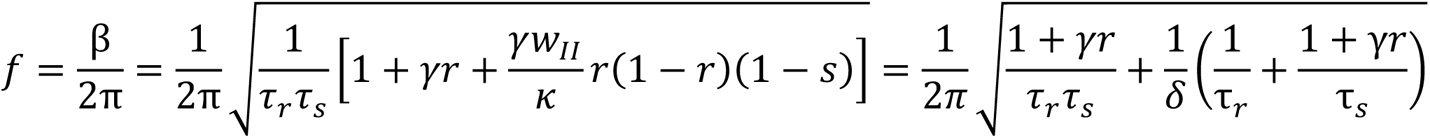

Therefore, as *δ* increases, the frequency *f* decreases.

**S7 Fig.**
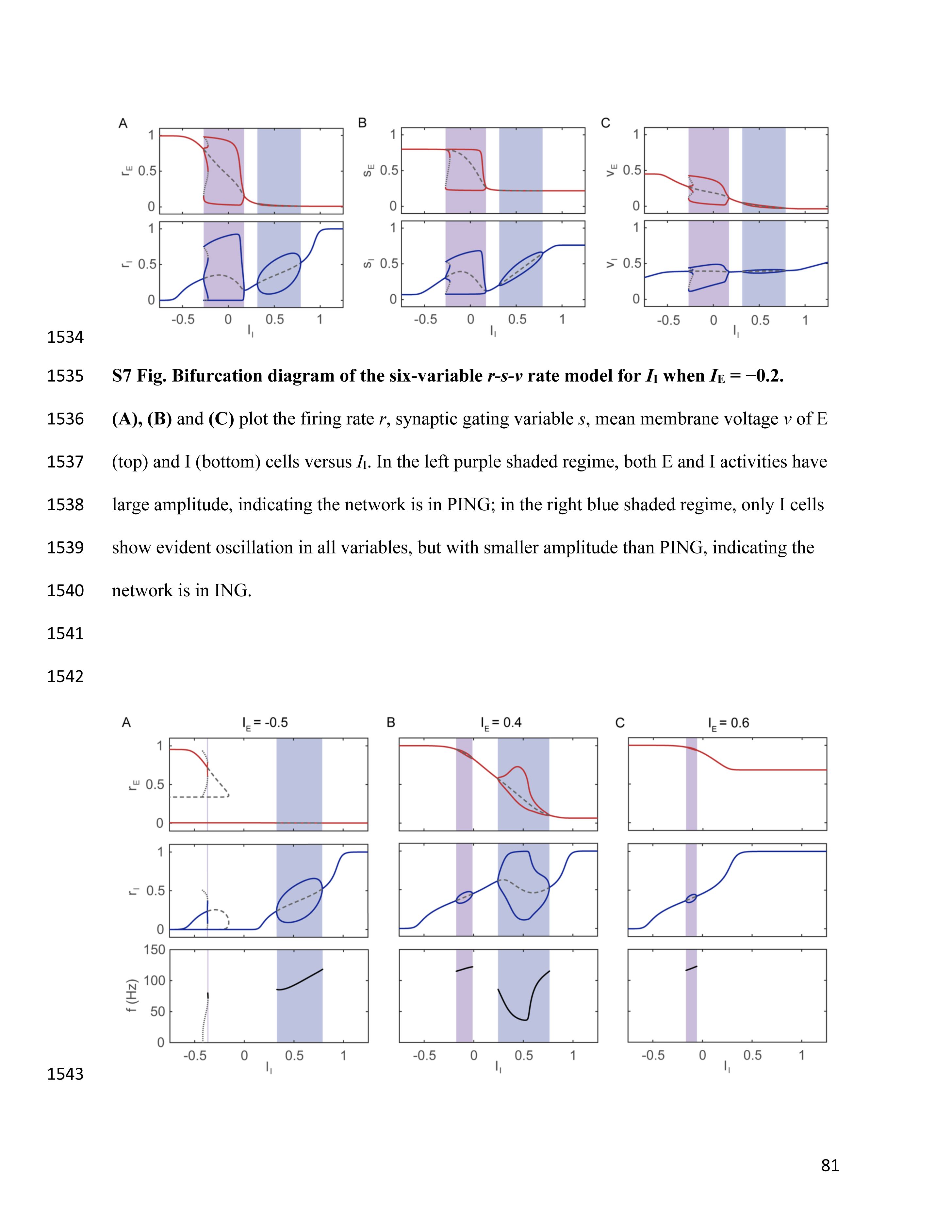
Bifurcation diagram of the six-variable *r-s-v* rate model for *I*_I_ when *I*_E_ = −0.2. **(A), (B)** and **(C)** plot the firing rate *r*, synaptic gating variable *s*, mean membrane voltage *v* of E (top) and I (bottom) cells versus *I*_I_. In the left purple shaded regime, both E and I activities have large amplitude, indicating the network is in PING; in the right blue shaded regime, only I cells show evident oscillation in all variables, but with smaller amplitude than PING, indicating the network is in ING.

**S8 Fig.**
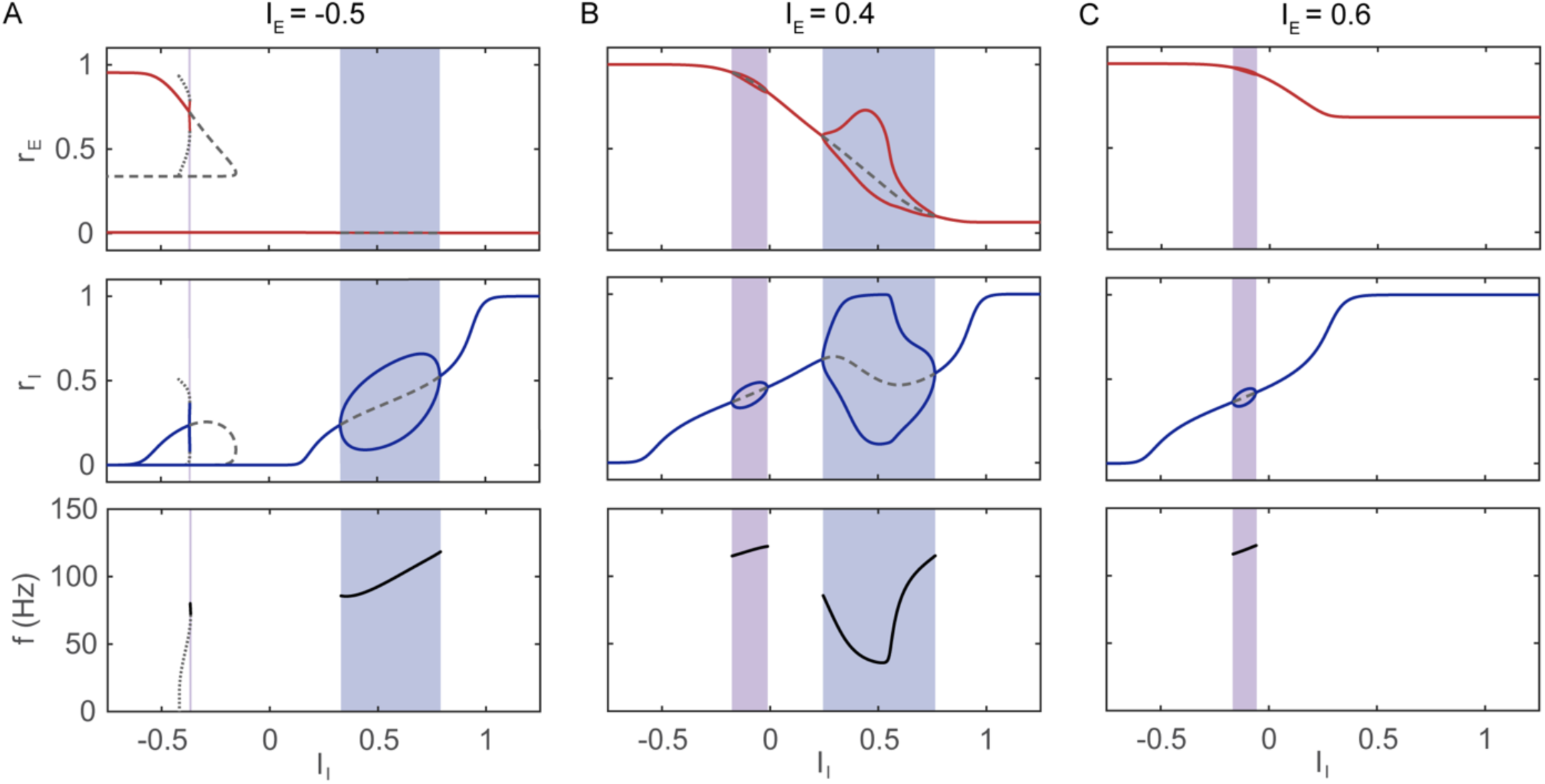
Bifurcation diagrams of the six-variable *r-s-v* rate model for small and large *I*_E_. **(A)** When *I*_E_ = −0.5, the PING regime (left) is destroyed because of the passage of a saddle-node bifurcation. The *r*_E_ - SSR has three intersections with the *r*_I_ - SSR, two of which are on the left flat branch of the *r*_I_ - SSR. **(B)** When *I*_E_ = 0.4, the oscillation amplitude in the left regime is tiny and does not correspond to either PING or ING; the right regime has a combination of PING and ING. **(C)** When *I*_E_ = 0.6, the right oscillation regime disappears due to the loss of Hopf bifurcation. There is no oscillation of biological interest when *I*_E_ is too large.

## Notes

### Competing Interest Statement

The authors have declared no competing interest.

## References

1. Bressler SL, Freeman WJ. Frequency analysis of olfactory system EEG in cat, rabbit, and rat. Electroencephalogr Clin Neurophysiol. 1980;50: 19–24.

2. Freeman WJ. Definitions of state variables and state space for brain-computer interface: Part 1. Multiple hierarchical levels of brain function. Cogn Neurodyn. 2007;1: 3–14. doi:10.1007/s11571-006-9001-x

3. Buzsáki G, Wang X-J. Mechanisms of gamma oscillations. Annu Rev Neurosci. 2012;35: 203–225. doi:10.1146/annurev-neuro-062111-150444

4. Penttonen M, Kamondi A, László A, Buzsáki G. Gamma frequency oscillation in the hippocampus of the rat: intracellular analysis in vivo. European Journal of Neuroscience. 1998;10: 718–728.

5. Mann EO, Radcliffe CA, Paulsen O. Hippocampal gamma-frequency oscillations: From interneurones to pyramidal cells, and back. Journal of Physiology. 2005;562: 55–63. doi:10.1113/jphysiol.2004.078758

6. Colgin LL, Moser EI. Gamma oscillations in the hippocampus. Physiology. 2010;25: 319–329. doi:10.1152/physiol.00021.2010

7. Chrobak JJ, Buzsá G. Gamma Oscillations in the Entorhinal Cortex of the Freely Behaving Rat. Journal of Neuroscience. 1998;18: 388–398.

8. Pinault D, Deschênes M. Voltage-dependent 40-Hz oscillations in rat reticular thalamic neurons in vivo. Neuroscience. 1992;51: 245–258.

9. Hughes JR. Responses from the visual cortex of unanesthetized monkeys. International review of neurobiology. Academic Press; 1964. pp. 99–152.

10. Adjamian P, Holliday IE, Barnes GR, Hillebrand A, Hadjipapas A, Singh KD. Induced visual illusions and gamma oscillations in human primary visual cortex. European Journal of Neuroscience. 2004;20: 587–592. doi:10.1111/j.1460-9568.2004.03495.x

11. Ray S, Maunsell JHR. Differences in Gamma Frequencies across Visual Cortex Restrict Their Possible Use in Computation. Neuron. 2010;67: 885–896. doi:10.1016/j.neuron.2010.08.004

12. Jia X, Tanabe S, Kohn A. Gamma and the Coordination of Spiking Activity in Early Visual Cortex. Neuron. 2013;77: 762–774. doi:10.1016/j.neuron.2012.12.036

13. Le Van Quyen M, Muller LE, Telenczuk B, Halgren E, Cash S, Hatsopoulos NG, et al. High-frequency oscillations in human and monkey neocortex during the wake-sleep cycle. Proc Natl Acad Sci U S A. 2016;113: 9363–9368. doi:10.1073/pnas.1523583113

14. Howard MW, Rizzuto DS, Caplan JB, Madsen JR, Lisman J, Aschenbrenner-Scheibe R, et al. Gamma Oscillations Correlate with Working Memory Load in Humans. Cerebral Cortex. 2003;13: 1369–1374. doi:10.1093/cercor/bhg084

15. Roux F, Uhlhaas PJ. Working memory and neural oscillations: Alpha-gamma versus theta-gamma codes for distinct WM information? Trends Cogn Sci. 2014;18: 16–25. doi:10.1016/j.tics.2013.10.010

16. Bauer M, Oostenveld R, Peeters M, Fries P. Tactile spatial attention enhances gamma-band activity in somatosensory cortex and reduces low-frequency activity in parieto-occipital areas. Journal of Neuroscience. 2006;26: 490–501. doi:10.1523/JNEUROSCI.5228-04.2006

17. Jensen O, Kaiser J, Lachaux J-P. Human gamma-frequency oscillations associated with attention and memory. Trends Neurosci. 2007;30: 317–324. doi:10.1016/j.tins.2007.05.001

18. Börgers C, Epstein S, Kopell NJ. Gamma oscillations mediate stimulus competition and attentional selection in a cortical network model. Proceedings of the National Academy of Sciences. 2008;105: 18023–18028.

19. Gray CM, König P, Engle Andreas K., Singer W. Oscillatory responses in cat visual cortex exhibit inter-columnar synchronization which reflects global stimulus properties. Nature. 1989;338: 334–337.

20. Lachaux JP, George N, Tallon-Baudry C, Martinerie J, Hugueville L, Minotti L, et al. The many faces of the gamma band response to complex visual stimuli. Neuroimage. 2005;25: 491–501. doi:10.1016/j.neuroimage.2004.11.052

21. Gross J, Schnitzler A, Timmermann L, Ploner M. Gamma oscillations in human primary somatosensory cortex reflect pain perception. PLoS Biol. 2007;5: 1168–1173. doi:10.1371/journal.pbio.0050133

22. Uhlhaas PJ, Singer W. Neural Synchrony in Brain Disorders: Relevance for Cognitive Dysfunctions and Pathophysiology. Neuron. 2006;52: 155–168. doi:10.1016/j.neuron.2006.09.020

23. Jefferys JGR, Traub RD, Whittington Miles A. Neuronal networks for induced ‘40 Hz’ rhythms. Trends Neurosci. 1996;19: 202–208.

24. Traub RD, Whittington MA, Stanford IM, Jefferys JGR. A mechanism for generation of long-range synchronous fast oscillations in the cortex. Nature. 1996;383: 621–624.

25. Börgers C, Kopell N. Synchronization in networks of excitatory and inhibitory neurons with sparse, random connectivity. Neural Comput. 2003;15: 509–538.

26. Whittington MA, Traubtt RD, Jefferys JGR. Synchronized oscillations in interneuron networks driven by metabotropic glutamate receptor activation. Nature. 1995;373: 612– 615.

27. Wang X-J, Buzsáki G. Gamma Oscillation by Synaptic Inhibition in a Hippocampal Interneuronal Network Model. Journal of neuroscience. 1996;16: 6402–6413.

28. Whittington MA, Traub RD, Kopell N, Ermentrout B, Buhl EH. Inhibition-based rhythms: experimental and mathematical observations on network dynamics. International Journal of Psychophysiology. 2000;38: 315–336.

29. Fisahn A, Pike FG, Buhl EH, Paulsen O. Cholinergic induction of network oscillations at 40 Hz in the hippocampus in vitro. Nature. 1998;394: 186–189.

30. Fisahn A, Contractor A, Traub RD, Buhl EH, Heinemann SF, McBain CJ. Distinct roles for the kainate receptor subunits GluR5 and GluR6 in kainate-induced hippocampal gamma oscillations. Journal of Neuroscience. 2004;24: 9658–9668. doi:10.1523/JNEUROSCI.2973-04.2004

31. Wilson HR, Cowan JD. Excitatory and Inhibitory Interactions in Localized Populations of Model Neurons. Biophys J. 1972;12: 1–24. doi:10.1016/S0006-3495(72)86068-5

32. Keeley S, Fenton AA, Rinzel J. Modeling fast and slow gamma oscillations with interneurons of different subtype. J Neurophysiol. 2017;117: 950–965. doi:10.1152/jn.00490.2016.-Experimental

33. Keeley S, Byrne Á, Fenton A, Rinzel J. Firing rate models for gamma oscillations. J Neurophysiol. 2019;121: 2181–2190. doi:10.1152/jn.00741.2018.-Gamma

34. Devalle F, Roxin A, Montbrió E. Firing rate equations require a spike synchrony mechanism to correctly describe fast oscillations in inhibitory networks. PLoS Comput Biol. 2017;13: e1005881. doi:10.1371/journal.pcbi.1005881

35. Börgers C, Walker B. Toggling between gamma-frequency activity and suppression of cell assemblies. Front Comput Neurosci. 2013;7: 33. doi:10.3389/fncom.2013.00033

36. Börgers C, Kopell N. Effects of Noisy Drive on Rhythms in Networks of Excitatory and Inhibitory Neurons. Neural Comput. 2005;17: 557–608.

37. Viriyopase A, Memmesheimer R-M, Gielen S. Cooperation and competition of gamma oscillation mechanisms. J Neurophysiol. 2016;116: 232–251. doi:10.1152/jn.00493.2015.-Os

38. Börgers C, Epstein S, Kopell NJ. Background gamma rhythmicity and attention in cortical local circuits: A computational study. Proceedings of the National Academy of Sciences. 2005;102: 7002–7007.

39. ter Wal M, Tiesinga P. Hippocampal Oscillations, Mechanisms (PING, ING, Sparse). In: Jaeger D, Jung R, editors. Encyclopedia of Computational Neuroscience. New York: Springer; 2013. pp. 1–14. doi:10.1007/978-1-4614-7320-6_475-3

40. Stiefel KM, Wespatat V, Gutkin B, Tennigkeit F, Singer W. Phase Dependent Sign Changes of GABAergic Synaptic Input Explored In-Silicio and In-Vitro. J Comput Neurosci. 2005;19: 71–85.

41. Chaudhuri R, Knoblauch K, Gariel MA, Kennedy H, Wang X-J. A Large-Scale Circuit Mechanism for Hierarchical Dynamical Processing in the Primate Cortex. Neuron. 2015;88: 419–431. doi:10.1016/j.neuron.2015.09.008

42. Brunel N, Wang XJ. What determines the frequency of fast network oscillations with irregular neural discharges? I. Synaptic dynamics and excitation-inhibition balance. J Neurophysiol. 2003;90: 415–430. doi:10.1152/jn.01095.2002

43. Honey CJ, Kötter R, Breakspear M, Sporns O. Network structure of cerebral cortex shapes functional connectivity on multiple time scales. Proceedings of the National Academy of Sciences. 2007;104: 10240–10245.

44. Byrne Á, O’Dea RD, Forrester M, Ross J, Coombes S. Next-generation neural mass and field modeling. J Neurophysiol. 2020;123: 726–742.

45. Bartos M, Vida I, Jonas P. Synaptic mechanisms of synchronized gamma oscillations in inhibitory interneuron networks. Nat Rev Neurosci. 2007;8: 45–56. doi:10.1038/nrn2044

46. Golomb D, Rinzel J. Dynamics of globally coupled inhibitory neurons with heterogeneity. Phys Rev E. 1993;48: 4810.

47. White JA, Chow CC, Ritt J, Soto-Treviño C, Kopell N. Synchronization and Oscillatory Dynamics in Heterogeneous, Mutually Inhibited Neurons. J Comput Neurosci. 1998;5: 5– 16.

48. Neltner L, Hansel D, Mato G, Meunier C. Synchrony in Heterogeneous Networks of Spiking Neurons. Neural Comput. 2000;12: 1607–1641.

49. Tiesinga PHE, José J V. Robust gamma oscillations in networks of inhibitory hippocampal interneurons. Network: Computation in Neural Systems. 2000;11: 1.

50. Tikidji-Hamburyan RA, Martínez JJ, White JA, Canavier CC. Resonant interneurons can increase robustness of gamma oscillations. Journal of Neuroscience. 2015;35: 15682– 15695. doi:10.1523/JNEUROSCI.2601-15.2015

51. Ernst U, Pawelzik K, Geisel T. Synchronization Induced by Temporal Delays in Pulse-Coupled Oscillators. Phys Rev Lett. 1995;74: 1570.

52. Maex R, De Schutter E. Resonant Synchronization in Heterogeneous Networks of Inhibitory Neurons. Journal of Neuroscience. 2003;23: 10503–10504.

53. Bartos M, Vida I, Frotscher M, Meyer A, Monyer H, Rg J, et al. Fast synaptic inhibition promotes synchronized gamma oscillations in hippocampal interneuron networks. Proceedings of the National Academy of Sciences. 2002;99: 13222–13227.

54. Gibson JR, Beierlein M, Connors BW. Functional properties of electrical synapses between inhibitory interneurons of neocortical layer 4. J Neurophysiol. 2005;93: 467–480. doi:10.1152/jn.00520.2004

55. Lewis TJ, Rinzel J. Dynamics of Spiking Neurons Connected by Both Inhibitory and Electrical Coupling. J Comput Neurosci. 2003;14: 283–309.

56. Vida I, Bartos M, Jonas P. Shunting inhibition improves robustness of gamma oscillations in hippocampal interneuron networks by homogenizing firing rates. Neuron. 2006;49: 107–117. doi:10.1016/j.neuron.2005.11.036

57. Tikidji-Hamburyan RA, Canavier CC. Shunting inhibition improves synchronization in heterogeneous inhibitory interneuronal networks with type 1 excitability whereas hyperpolarizing inhibition is better for type 2 excitability. eNeuro. 2020;7. doi:10.1523/ENEURO.0464-19.2020

58. Brunel N, Hakim V. Sparsely synchronized neuronal oscillations. Chaos: An Interdisciplinary Journal of Nonlinear Science. 2008;18: 015113. doi:10.1063/1.2779858

59. Brunel N, Hakim V. Fast Global Oscillations in Networks of Integrate-and-Fire Neurons with Low Firing Rates. Neural Comput. 1999;11: 1621–1671.

60. Tikidji-Hamburyan RA, Leonik CA, Canavier CC. Phase response theory explains cluster formation in sparsely but strongly connected inhibitory neural networks and effects of jitter due to sparse connectivity. J Neurophysiol. 2019;121: 1125–1142. doi:10.1152/jn.00728.2018

61. Van Vreeswijk C, Abbott LF, Ermentrout GB. When Inhibition not Excitation Synchronizes Neural Firing. J Comput Neurosci. 1994;1: 313–321.

62. Geisler C, Brunel N, Wang XJ. Contributions of intrinsic membrane dynamics to fast network oscillations with irregular neuronal discharges. J Neurophysiol. 2005;94: 4344– 4361. doi:10.1152/jn.00510.2004

63. Schieferstein N, Schwalger T, Lindner B, Kempter R. Intra-ripple frequency accommodation in an inhibitory network model for hippocampal ripple oscillations. BioRxiv 2023.01.30.526209 [Preprint]. 2023 [cited 2023 Apr 27]. Available from: https://www.biorxiv.org/content/10.1101/2023.01.30.526209v1 doi:10.1101/2023.01.30.526209

64. Ahmadian Y, Rubin DB, Miller KD. Analysis of the stabilized supralinear network. Neural Comput. 2013;25: 1994–2037. doi:10.1162/NECO_a_00472

65. Jadi MP, Sejnowski TJ. Regulating cortical oscillations in an inhibition-stabilized network. Proceedings of the IEEE. 2014;102: 830–842. doi:10.1109/JPROC.2014.2313113

66. Tsodyks M V, Skaggs WE, Sejnowski TJ, Mcnaughton BL. Paradoxical Effects of External Modulation of Inhibitory Interneurons. Journal of neuroscience. 1997;17: 4382– 4388.

67. Kopell N, Börgers C, Pervouchine D., Malerba P., Tort A. Gamma and theta rhythms in biophysical models of hippocampal circuits. Hippocampal Microcircuits. 2010;5: 423–457. doi:10.1007/978-1-4419-0996-1

68. Tabak J, Senn W, O’Donovan MJ, Rinzel J. Modeling of Spontaneous Activity in Developing Spinal Cord Using Activity-Dependent Depression in an Excitatory Network. The Journal of Neuroscience. 2000;20: 3041–3056.

69. Ermentrout B. Simulating, Analyzing, and Animating Dynamical Systems. Philadelphia: Society for Industrial and Applied Mathematics; 2002. doi:10.1137/1.9780898718195

70. Hsu T-H. Available from: https://sites.pitt.edu/~phase/bard/bardware/xpp/plotxppaut4p4.m.

71. Strang G. Introduction to linear algebra. 5th ed. Wellesley: Wellesley - Cambridge Press; 2016.

